# Scanning and active sampling behaviours emerge from conserved insect neural circuits

**DOI:** 10.1101/2025.10.13.682010

**Authors:** Cody A Freas, Antoine Wystrach

**Author notes:** Corresponding Author: Address for correspondence: Research Center on Animal Cognition (CRCA) Center for Integrative Biology (CBI), CNRS, University of Toulouse, Toulouse, France.

## Abstract

Navigating insects often pause and rotate to sample their surroundings, behaviours termed scanning. These and other active sampling behaviours embody navigational uncertainty, and are key for spatial learning, yet their neural basis remains unclear and existing models impose scanning behaviours rather than explaining its emergence. Here, we show that desert ants’ scanning dynamics can emerge spontaneously from the same conserved neural circuits used for goal-directed navigation, without requiring a specialized scanning module. We built a biologically grounded model combining central complex (CX) steering and lateral accessory lobe (LAL) oscillators, and added a downstream stochastic inhibition of forward speed. This minimal system produced diverse, realistic scan dynamics; saccades, fixations and reversals, whose features were qualitatively compared to high-speed video recordings of *Melophorus bagoti* scanning. Detailed analysis of these natural scans confirmed model predictions, including how scan structure depends on oscillator phase, goal-heading deviation, and navigational uncertainty. Furthermore, the model reveals that simple modulation of forward speed unifies a broad range of behaviours across ant species, from dashes to smooth oscillatory trajectories to pirouettes and voltes. Crucially, this model suggests a distributed control principle where forward speed acts as a single adjustable parameter, for both individuals and through evolution, to regulate the balance between goal-driven exploitation and information-seeking exploration.

## Background

Insects will often interrupt their forward movement and rotate to sample visual information from multiple body orientations. These behaviours appear widespread across both walking, crawling and flying insects (as well as other taxa), and take various forms and names such as ‘scans’, ‘volte’, ‘pirouette’, ‘turn back’ ‘peering’, ‘sweep’ or ‘dance’ (Baird et al., 2012; Gomez-Marin and Louis, 2014; Deeti et al., 2023a; Fleischmann et al., 2017; Graham and Collett, 2002; Lehrer, 1993; Mouritsen et al., 2004; Müller and Wehner, 2010; Stürzl et al., 2016; Tarsitano and Andrew, 1999; Ugolini, 2006; Wallace, 1959; Wehner et al., 1992; Wystrach et al., 2014; Zeil and Fleischmann, 2019). Despite its prevalence across taxa, the neural mechanisms underlying these seemingly highly structured behaviours remain unclear.

Scans are especially prevalent in visually guided ants (Deeti et al., 2023a; Fleischmann et al., 2017; Freas et al., 2018; Müller and Wehner, 2010; Nicholson et al., 1999; Wehner et al., 1996; Wystrach et al., 2014; Zeil and Fleischmann, 2019). Ants occasionally cease forward motion, and the ensuing scanning behaviours consist of distinct periods: saccades, when the navigator is rotating, and fixations, when the ant pauses all movement (Figure 1A-C; Video 1,2, Müller and Wehner, 2010; Wystrach *et al*., 2014; Zeil and Fleischmann, 2019; Deeti, Cheng, *et al*., 2023; Freas and Cheng, 2025). Multiple saccades and fixations make up a singular scan, which may contain one or more rotational reversals (right then left/left then right) before forward movement resumes. The probability of scanning, while stochastic (Deeti et al., 2023a), increases with uncertainty, when views are unfamiliar or unexpected (Deeti et al., 2023a; Freas et al., 2018; Schwarz et al., 2020b; Wystrach et al., 2014) such as during learning walks (Müller and Wehner, 2010; Zeil and Fleischmann, 2019) and novel route formation (Freas and Cheng, 2025, 2022; Haalck et al., 2023), or when views have been associated with aversive events (Freas et al., 2022; Wystrach et al., 2020a).

**Figure 1.**
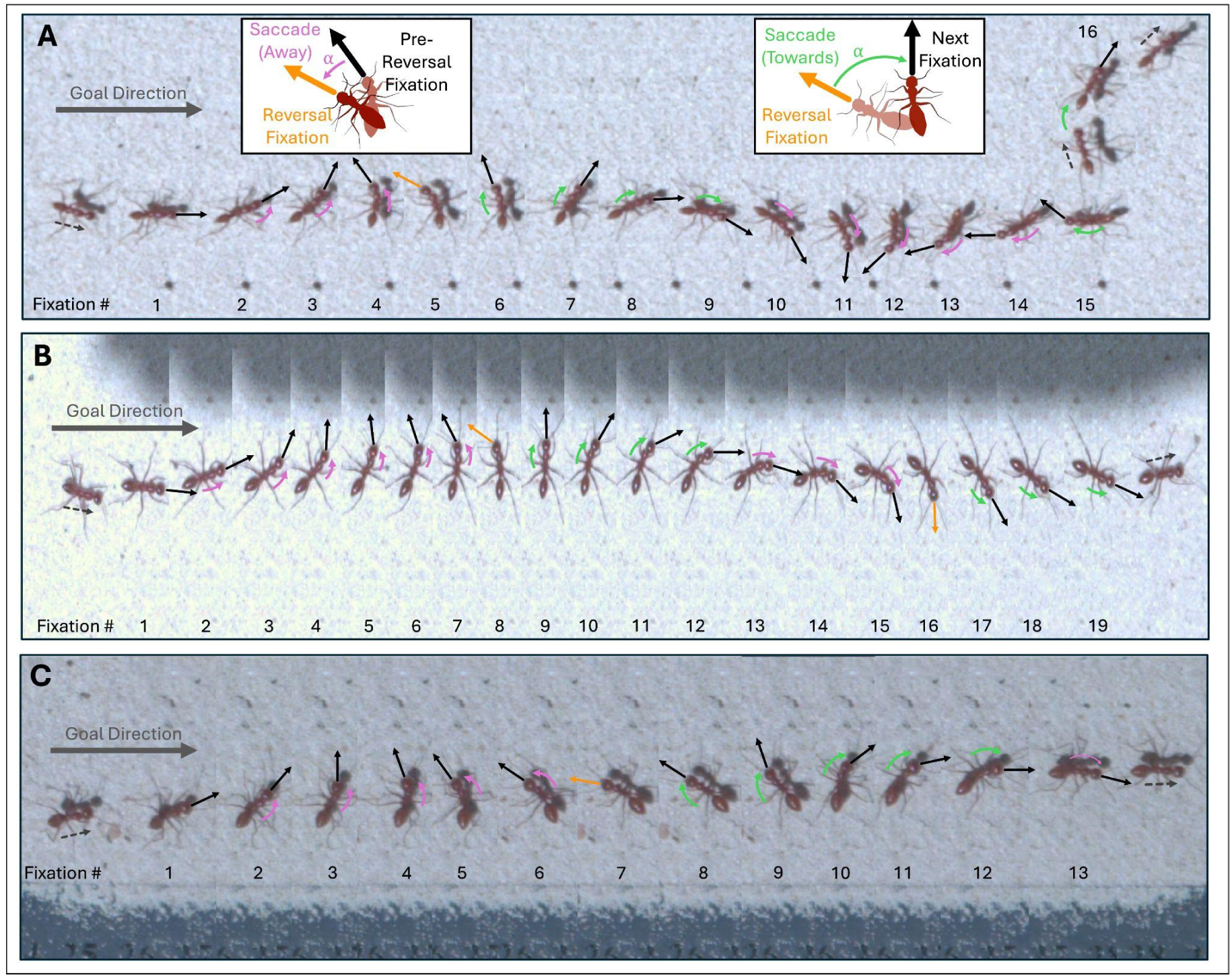
Image composites of fixations (pauses – typically 50-150ms) during three different examples of scanning behaviours in *Melophorus bagoti*, all taken within 1m of the nest entrance, in inexperienced ants forming their route to a feeder. Images were extracted and compiled from highspeed video taken at 600fps at 1080p using a Chronos 2.1HD camera (field of view, 30 cm × 17 cm). For each image, fixations are indicated by consecutively numbered black arrows denoting the ant’s orientation (except reversal). Rotational movement between fixations, saccades, are classified as either away (pink) or towards (green) the goal direction (∼90°). When a fixation precedes a change in turning (from left to right in these examples), this fixation is classified as a reversal (orange). (**A**) Denotes a scanning example where it reverses direction once but then turns in a complete loop with no reversal (fixations 6-16). (**B**) Shows a scan which contains multiple reversals (two). (**C**) Illustrates a scan with one reversal, ∼180° from the goal direction. Black dotted arrows denote pre/post scan forward movement.

Functionally, scans are associated with information sampling, helping ants choose headings when cues are unfamiliar or in conflict (Wystrach et al., 2014; Wystrach, Buehlmann, *et al*., 2020; Deeti, Cheng, *et al*., 2023; Freas and Cheng, 2025). Additionally, their prevalence during both learning walks and early route formation indicates a role in view acquisition (Freas and Cheng, 2025; Müller and Wehner, 2010; Wystrach, 2023; Wystrach et al., 2014; Zeil, 2023; Zeil and Fleischmann, 2019).

Our mechanistic understanding of insect navigation has improved tremendously in the last decades, thanks to a combination of behavioural, neurobiological and modelling studies (Honkanen et al., 2019; Webb and Wystrach, 2016). However, current models do not explain the emergence and natural dynamics of scanning behaviours. Instead, ‘scanning routines’ are sometimes force-added in models to help the agent or robot sample views across directions (Baddeley et al., 2012; Gattaux et al., 2023; Husbands et al., 2021; Knight et al., 2019; Le Möel and Wystrach, 2020; Murray et al., 2019; Wystrach et al., 2013).

Here we ask whether and how ants’ scanning dynamics could emerge from neural structures typically implicated in insect navigation: the central complex (CX) and the lateral accessory lobes (LAL). While the circuitry of these structures is detailed primarily in a few insect model species, it is sufficiently conserved to allow for neural modelling of insect navigation in various contexts (Adden et al., 2022; Clément et al., 2023; Collett et al., 2025; Goulard et al., 2023; Shiu et al., 2024; Steinbeck et al., 2020; Stone et al., 2017; Webb and Wystrach, 2016; Wystrach et al., 2020b). Both CX and LAL regions are well established to integrate multiple directional information and control steering commands in navigating insects (Franconville et al., 2018; Heinze, 2017; Honkanen et al., 2019; Hulse et al., 2021; Li et al., 2020; Namiki and Kanzaki, 2016; Pfeiffer and Homberg, 2014).

Previous work has established that goal headings are encoded in the central complex (CX) (Mussels Pires et al., 2024) and modelling efforts have shown that this circuitry can naturally accommodate innate and learnt guidance such as path integration, learn vectors, visual route following or homing as observed in ants and bees (Goulard et al., 2021; Honkanen et al., 2019; Stone et al., 2017; Le Moël et al. 2019; Wystrach, 2023; Wystrach et al., 2020b). In parallel, oscillatory dynamics in the lateral accessory lobes (LAL), produced by reciprocal inhibition across both hemispheres and conveyed by so-called descending flip-flopping neurons, were shown to drive the spontaneous zigzags displayed by moths upon losing their pheromone plume (Kanzaki and Mishima, 1996; Mishima and Kanzaki, 1998, 1999; Wada and Kanzaki, 2005; Kanzaki et al., 2005; Iwano et al., 2010). Subsequent modelling efforts have shown how these circuits can equally support the continuous lateral oscillations displayed by a wide range of insect species (Adden et al., 2022; Kanzaki et al., 2005; Namiki and Kanzaki, 2016; Steinbeck et al., 2020; Wystrach et al., 2016), including ants (Clément et al., 2023; Dauzere-Peres et al., 2026). Building on these elements, the present work addresses a distinct question that current navigation models do not resolve: whether discrete active sampling behaviours such as ant scanning require specialised control mechanisms, or instead emerge from interactions within these conserved navigation circuits.

To investigate this, we analysed high-speed recordings of natural scanning behaviours in *Melophorus bagoti* foragers and asked whether their detailed dynamics could be captured by modelling the interactions between the CX and the LAL, under the condition of occasional halted forward movement. We developed a biologically constrained neural model of a CX steering output that modulates a downstream LAL oscillator, under the assumption that forward speed may be transiently gated by a stochastic extrinsic signal (Deeti et al., 2023). Finally, we added the physical constraint that forward velocity increasingly limits rotation. Without introducing a dedicated scanning module, this model is sufficient to reproduce multiple key qualitative features of natural scans, including the relative durations and amplitudes of saccades, fixations, directional reversals, and the emergence of rare full loop scans, which arise as a consequence of CX–LAL and forward-angular speed interactions.

In addition to behaviours typically categorized as ‘scans,’ the model produces information sampling behaviours often described using different terminology; including ‘pirouettes’ observed in ants (Fleischmann et al., 2017; Zeil and Fleischmann, 2019), dung beetle ‘dances’ (Baird et al., 2012), and *Drosophila* ‘reorientations’ (Demir et al., 2020; Gomez-Marin et al., 2011). Furthermore, by reducing rather than terminating forward speed, the model also captures continuous-movement behaviours such as voltes and large oscillating paths seen across ant species (Fleischmann et al., 2017; Zeil and Fleischmann, 2019; Clément, Schwarz and Wystrach, 2023). These findings point to a conserved neural control strategy where forward speed inhibition gates the expression of angular motor output, allowing for shifting seamlessly between information gathering and goal-oriented progression. The few novel assumptions added here enable us to form predictions for future work.

### Neural substrates of insect navigation

To interpret these results, we briefly summarise the neural substrates that motivated the model. The CX tracks the insects’ heading relative to the world, transforms egocentric sensory information into allocentric current and goals directions, and compares both to generate steering commands. Within the CX’s substructures, the current heading direction is represented neurally in the protocerebral bridge (PB) and the Ellipsoid Body (EB) (Kim et al., 2019; Seelig and Jayaraman, 2015; Honkanen et al., 2019; Pfeiffer and Homberg, 2014), while goal directions have been shown to be represented in the fan-shaped body (FB) (Mussells Pires et al., 2024), consistent with earlier modelling (Stone et al., 2017). Current and goal direction representations are compared within the CX and project left/right steering commands to the LAL (Green et al., 2019; Honkanen et al., 2019; Iwano et al., 2010; Mussells Pires et al., 2024; Westeinde et al., 2024; Figure 2A).

**Figure 2.**
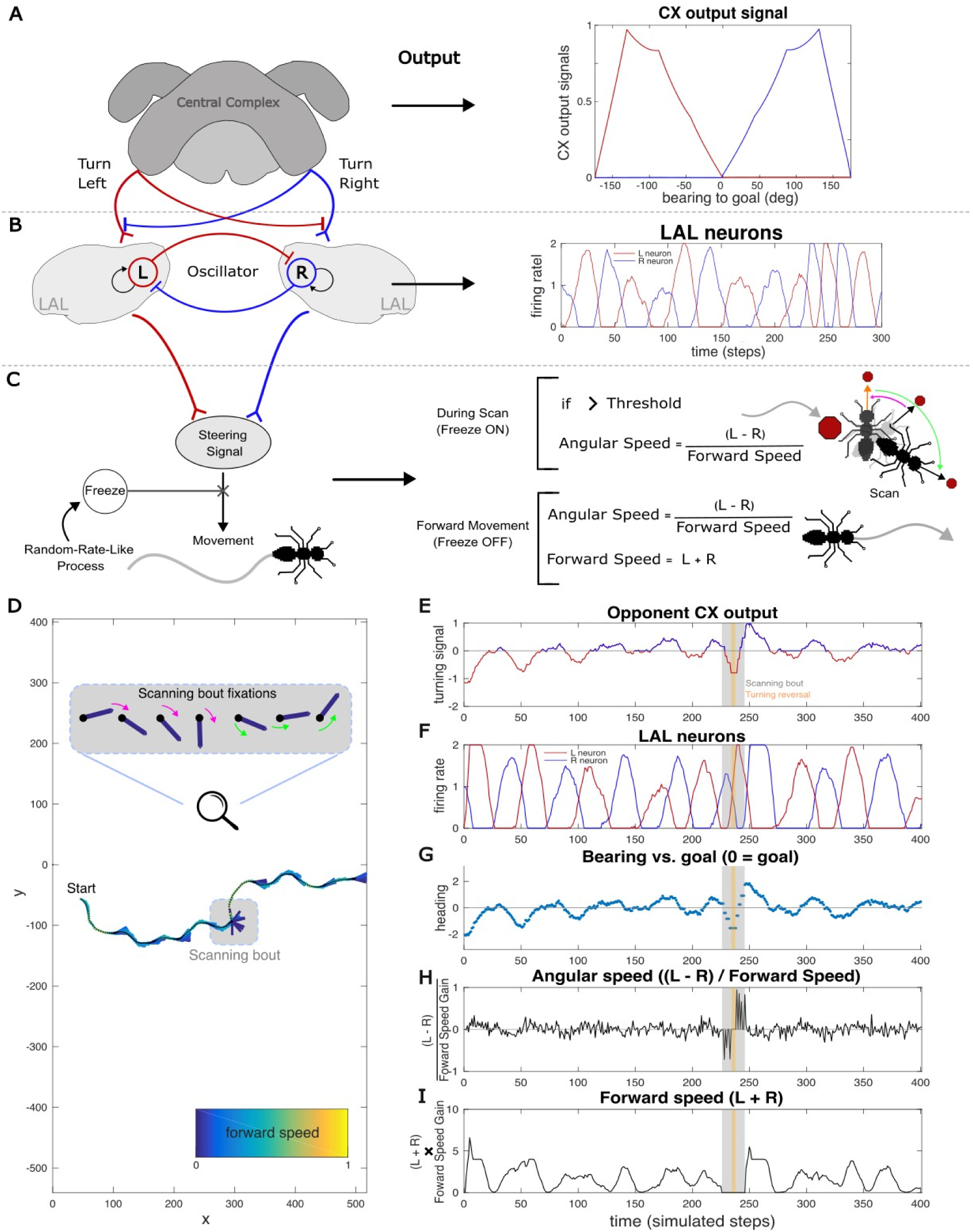
Schematic of neural circuit model and its outputs across navigation relevant brain regions. (**A**) The central complex (CX), a mid-brain region whose subregions compare representations of the goal heading with a representation of the agent’s compass-based current heading. The CX outputs bilateral signals to turn the left (red line) and right (blue line) to the Lateral Accessory Lobes (LAL). (**B**) Within the LAL oscillator, left (L) and right (R) neurons reciprocally inhibit one another (red and blue connections), while attempting to maintain a basal firing rate via internal feedback (circular black arrows), forming an oscillator which outputs a stable anti-phasic oscillatory activity between the L and R neurons across time (red and blue lines). (**C**) A steering signal is outputted from the LAL and results in the agent’s forward and angular movement. This steering signal stopped via an external inhibitory ‘freeze signal’, that breaks the angular and forward speed central pattern generators (CPG), initiating the scan. A threshold was implemented for the underlying angular drive to restart the CPG after each fixation period, resulting in a saccade whose magnitude was determined by this accumulated angular drive. The agent’s angular speed was normalised by the forward speed (linear normalisation - angular speed = angular drive/(forward speed + 0.1)) so the agent produces larger saccades magnitudes during scans. Threshold was determined to roughly mirror saccade magnitudes in real-world ants. Scan duration was implemented as an exponential duration distribution through a dice-roll at each simulation step. (**D**) Characteristics of an example path, with a single scanning bout, generated by the model. Black dots represent the agent’s head position at each simulated step coupled with coloured bars which indicate both the forward speed (arbitrary scale) and heading direction of each step. During the scanning bout, the agent’s forward speed is zero and the model produces several fixations in separate directions. The fixations of this scanning bout are zoomed and separated into a sequence of fixations. Coloured arrows indicate saccades, rotational movements between fixations which were defined as either away (pink) or towards (green) the goal direction. (**E**) The modelled CX’s turning signal output towards the goal direction, based on if the agent’s heading direction is to the left or right of the goal. **(F)** The oscillatory activity between the R and L neurons in the LAL during the example path. The agent’s (**G**) heading direction, (**H**) angular speed, and (**I**) forward at each step during the path.

The CX receives convergent input from multiple sensory pathways (Honkanen et al., 2019) while behavioural and modeling evidence shows that ants combine cues such as wind and odours (Buehlmann et al., 2020; Knaden and Graham, 2016; Müller and Wehner, 2007; Steck et al., 2011), visual familiarity (computationally linked to Mushroom body output: Wystrach et al., 2020b), and path integration (modelled as a CX computation: Lu et al., 2022; Lyu et al., 2022; Stone et al., 2017) to set a goal direction. While the neural locus of this integration has not been directly recorded across all these modalities, the convergence architecture of the CX (Honkanen et al., 2019) makes it the most plausible substrate, and we assume such a goal direction exists in the FB of the CX.The LAL are premotor centers that act as a bottleneck, integrating information from multiple higher processing centres and sensory inputs, including outputs from the CX, and relay these signals via descending neurons that transmit motor commands to the thorax (Namiki and Kanzaki, 2016; Shih et al., 2015; Steinbeck et al., 2020). The LAL also produce long-lasting, alternation between left and right turn via so-called flip-flop neurons (Berni, 2015; Iwano et al., 2010; Kanzaki, 2005; Namiki and Kanzaki, 2016; Namiki et al., 2014, Steinbeck et al., 2020), which can result in the regular lateral oscillations observed in many insects (Freas and Cheng, 2022; Izquierdo and Lockery, 2010; Kanzaki et al., 1992; Kuenen and Baker, 1983; Namiki and Kanzaki, 2016; Olberg, 1983; Wystrach et al., 2016), notably in ants (See Video 4, Clément et al., 2023; Collet et al., 2025; Deeti and Cheng, 2025; Haalck et al., 2023; Hangartner, 1967). The generation of oscillatory movements allows for continuous spatial sampling of information, and modeling studies demonstrate that when these oscillations are modulated by odor or visual cues, they offer a highly effective strategy for odour plume or gradient tracking, visual route following or goal pinpointing (Adden et al., 2022; Kodzhabashev and Mangan, 2015; Le Möel and Wystrach, 2020; Wystrach et al., 2016).

Together, this CX–LAL circuitry is well suited to control continuous steering and sampling during locomotion (Collett et al., 2025). However, until now it is unknown if this same circuitry underlies the discrete structures of ant scanning behaviour without invoking a dedicated control routine.

## Results and Discussion

### A simple model for scanning behaviour

We first constructed a simple model of the insect central complex (CX; Figure 2A), following standard approaches (Goulard et al., 2023; Stone et al., 2017; Wystrach, 2023; Wystrach et al., 2020b). This circuit compares the agent’s current heading to a stored goal heading (which, in our simulations, is directionally fixed) and generates lateralised steering signals to reduce heading error. Unlike previous models (but as in Adden *et al*., 2022) the CX output here does not steer the agent directly, but instead modulates an intrinsic oscillator that generates alternating turns (Figure 2B), a mechanism consistent with the continuous oscillatory patterns known to underlie ant navigation (Clément, Schwarz and Wystrach, 2023).

Oscillatory turning behaviours are widespread in insects(Clément, Schwarz and Wystrach, 2023; Cheng, 2024) and underlie models of insect steering (Kanzaki, 2005; Kanzaki and Mishima, 1996). We modelled this as an intrinsic oscillator located in the lateral accessory lobes (LAL), a pre-motor region that receives, among others, CX outputs (Heinze, 2017) and is a proposed substrate for steering control (Steinbeck, Adden and Graham, 2020; Clément, Schwarz and Wystrach, 2023). The difference between these left and right signals, such as conveyed by descending neurons (DN) to the thoracic ganglia (Büschges and Ache, 2025), determines the agent’s angular speed, thereby generating the rhythmic turning dynamics akin to those observed in ants (Clément, Schwarz and Wystrach, 2023). In combination with the CX modulation, this oscillator produces a continuous oscillatory trajectory generally oriented towards the goal direction set in the CX (which is always to the right in our examples; Figure 2D).

To generate scanning behaviour (Figure 1A–C), we added a mechanism to intermittently interrupt forward movement, as observed in desert ants, where pauses occur randomly (Deeti et al., 2023a). Because we did not want to form assumptions for how such a ‘freeze signal’ could be implemented in the insect nervous system; in our model this was achieved using a simple external signal that halts forward motion at random intervals. Since scan duration in real ants follows a Poisson-like distribution (Deeti et al., 2023a), we approximated this by performing at each simulation step a dice-roll with equal probability (calibrated as 1/mean of ants observed scan duration) to restart forward motion, producing the desired Poisson distribution of scans’ duration.

During scanning, real ants display rotational saccades of variable duration and angular magnitude (Figure 1A–C). To replicate this, we introduced a threshold-based mechanism: after each fixation (i.e., zero angular and forward speed), the underlying angular steering signal accumulates until surpassing a threshold, triggering a saccade. The resulting angular magnitude of the saccade corresponds to the sum of the angular drive accumulated during the fixation. Here also we stuck to a non-neural, straightforward algorithmic level, as we did not want to make assumptions about how such a cumulate-and-release mechanism could be neurally implemented in the insect brain (see discussion for potential implementations).

With this, the agent now scans, but the heading deviation during scanning remains within the same limited range as during forward movement, highly constrained towards the goal direction. This does not reflect real scans, which involve much larger angular deviations (Deeti et al., 2023a; Fleischmann et al., 2017; Freas et al., 2019; Freas and Cheng, 2025; Müller and Wehner, 2010; Wystrach et al., 2014). We hypothesized that this limited turning during forward movement is due in part to biomechanical constraints: fast forward motion impedes sharp turns, while slowing down allows easier rotation. When ants stop, they are free to execute sharp turns via leg coordination that would be otherwise impossible (e.g., inner legs moving backward and outer legs forward), enabling high rotation during scans (Figure 1). To implement this constraint, we normalized the agent’s angular speed by the inverse of forward speed: the slower the agent moves, the more it turns for a given steering signal. As a result, the agent now produces larger saccades and greater heading deviations during scans, while remaining more constrained during forward movement. To match the saccade amplitudes observed in real ants, we adjusted the threshold for breaking fixation so that the resulting distribution of saccade angles (Figure 3A) approximates those seen in ants (Figure 3B). Higher thresholds yield less frequent but larger saccades.

**Figure 3.**
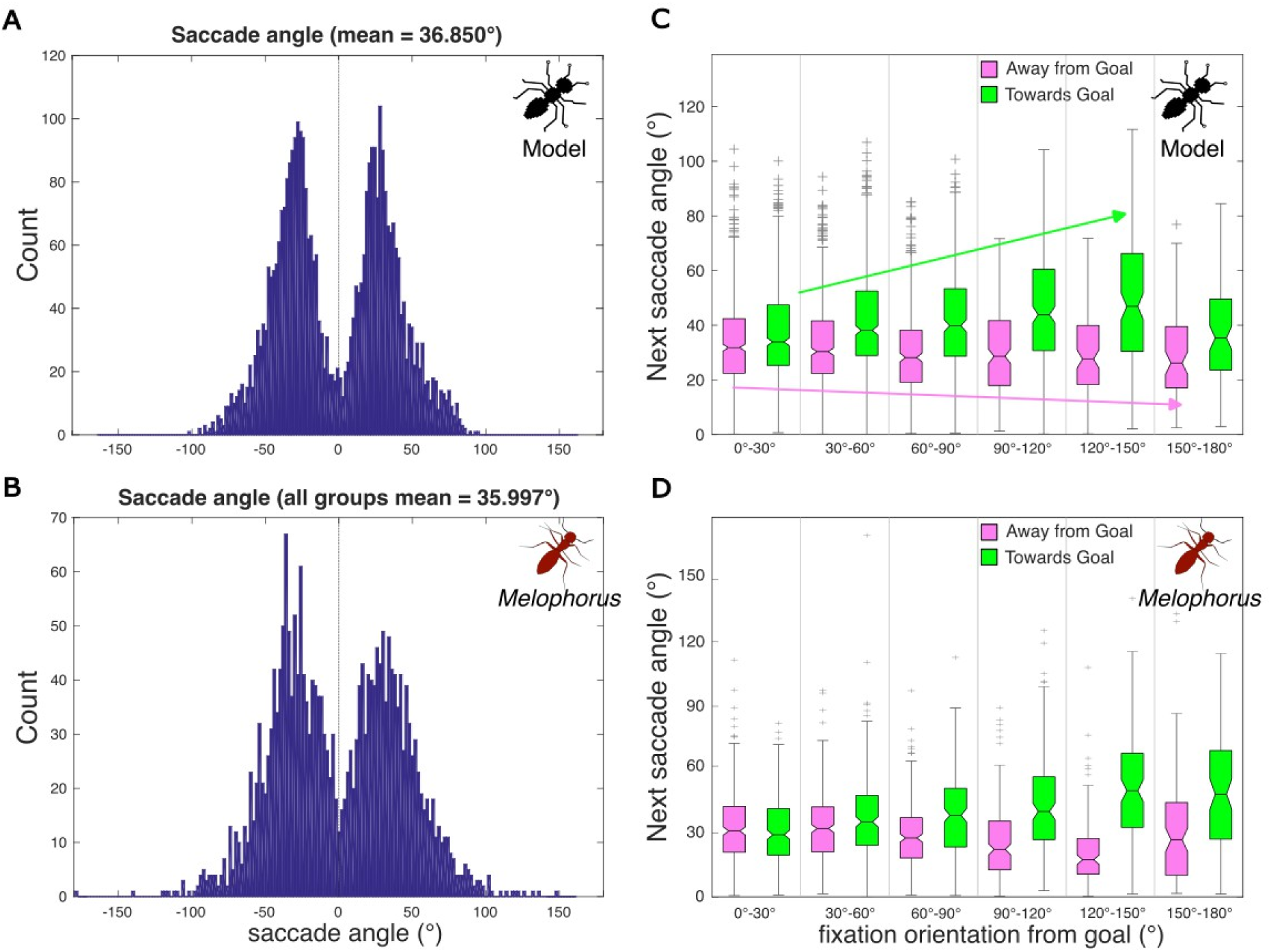
Saccade angle distribution and the effect of CX steering guidance on saccade angle magnitude. Saccade angle distributions in (**A**) the modelled agent and (**B**) real ants with all scan conditions combined. After each fixation, the subsequent saccade angle is plotted against the fixation’s angular divergence from the goal direction in both (**C**) the model agent and (**D**) real ants with all conditions combined. The general trends of the model are classified through coloured arrows (away-green, towards-pink) representing the increasing next saccade angle towards the goal direction and the decreasing next saccade angle away from the goal direction of the modelled agent when fixation orientation from the goal direction was large. This pattern is consistent with the model assumption that CX steering strength varies with angular divergence from the goal; easier to turn towards goal and thus large saccades, while harder to turn further away and small saccades. A similar pattern is observed in the ant data. For the box and whisker plots in panels C and D, the box spans the interquartile range while the horizontal line indicates the mean. Whiskers extend to the IQR X 1.5 while outliers beyond this range are shown as ‘+’ symbols. The indentation of the bar around the mean indicates the 95% confidence interval.

Overall, the interaction between the CX output (Figure 2A), oscillator phase (Figure 2B), and the scan control mechanism (Figure 2C) now produces complex, nonlinear dynamics, giving rise to emergent behaviours, which we explored and compared to real ants in the following subsections.

### CX steering influences saccade magnitude

A key prediction of our model is that the CX steering mechanism operates continuously, including during scans, and its influence should be detectable in the scan’s structure. Specifically, the strength of the CX’s corrective signal towards the goal increases with the agent’s angular deviation from that goal (Figure 2A).

If guided solely by the CX, the agent would barely turn away from the goal but due to noise. However, actual steering is governed by the downstream oscillator in the LAL, which ultimately determines the direction of each saccade during scans. Saccades towards the goal occur when the oscillator is in phase with the CX corrective signal, resulting in stronger angular drive and therefore larger saccades. Conversely, saccades away from the goal arise when the oscillator and CX corrective signals are in conflict, the latter restraining the former, producing weaker angular drive and thus smaller saccades. Since the CX’s corrective signal roughly scales with angular deviation (Figure 2A), this difference in saccade amplitude (towards vs. away from goal) should be minimal when the agent is facing the goal direction but becomes more pronounced as it is facing away (Figure 3C).

This predicted pattern closely matches real ant data (Figure 3D). Across all datasets, including inexperienced and experienced ants, whether homing or foraging (treated as random error in our LME), saccade amplitude was significantly explained by both the saccade direction (towards and away from the goal) (*F*_(1,1890)_ = 5.08, *p* = 0.024) and angular deviation (*F_(_*_1,1890)_ = 18.69, *p* < 0.001) and, importantly, by their interaction (*F*_(1,1890)_ = 43.04, *p* < 0.001). Specifically, saccades towards the goal were larger than those turning away from it, and this difference increased as the ants faced away from their goal, supporting the model’s prediction. Together, this supports that both the oscillator and CX are at play during scanning, with the latter continuously pulling the ants towards its goal heading.

Our statistical analysis on saccade amplitude also revealed significant fixed effects for the individuals (LRstat = 200.25; p < 0.001) and conditions (LRstat = 49.08, p < 0.001) (see Statistical Methods for details) as well as an interaction between heading orientation (*angular deviation* from goal) and whether the previous fixation was a reversal (*F*_(1,1890)_ = 6.86, p = 0.009; Figure 4A), which we will address in the next sections.

**Figure 4.**
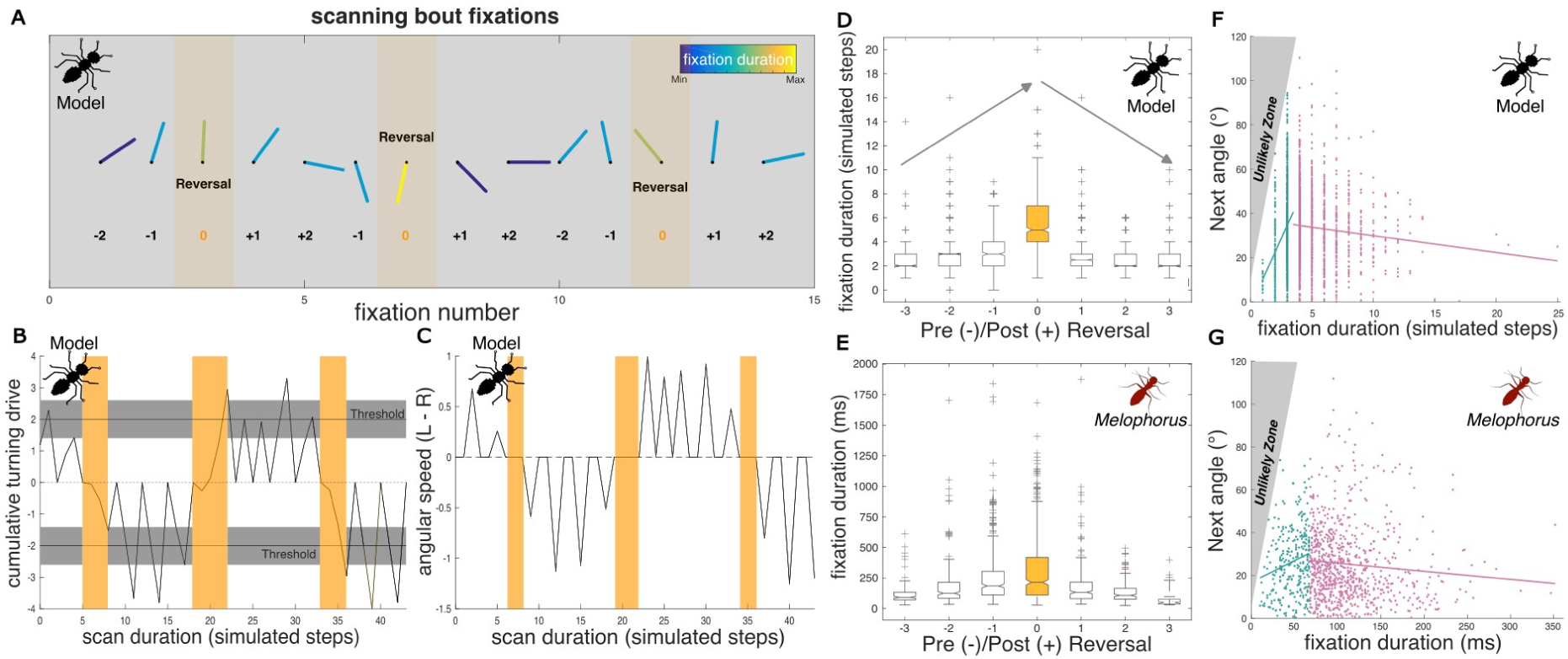
Relationship between fixation duration and reversal. (**A**) Sequence of fixation directions and their duration for an example scan containing multiple reversals (orange), showcasing that the longest fixations tend to occur when the ant reverses directions on the next saccade (fixation duration continuum, blue-min; yellow-max). (**B**) shows changes in the cumulative turning drive over the course of the example scan. To initiate a saccade, the drive must surpass a threshold (line). This threshold possesses a level of noise which leads to variance in when a saccade appears as well as to the variance in saccade magnitude. Reversals occur when the oscillatory cycle changes phase (±), which are associated with longer times for turning drive to accumulate beyond threshold (orange). (**C**) Angular speed through the simulation, spikes represent saccades to the left and right and the change from positive to negative illustrates reversal periods (orange). Relationship between fixation duration and oscillatory cycle changes (orange) and subsequently reversals, occur in (**D**) modelled and (**E**) real ants. In both, the longest fixation duration typically occurs at the reversal (0), increasing as the reversal approaches (-), while decreasing post reversal (+). For these box and whisker plots, the box spans the inter quartile range while the horizontal line indicates the mean. Whiskers extend to the IQR X 1.5 while outliers beyond this range are shown as ‘+’ symbols. The indentation of the bar around the mean indicates the 95% confidence interval. Correlation between fixation duration and the next saccade angle in (**F**) modelled and (**G**) real ants. Data are split into the lower 25% (Q1, Blue) and upper 75% (Q2-Q4, Pink) of the distribution. In both, the general tendency in Q1 is significantly positive (real ant; Linear regression model; F(1, 281) = 11.00, p = 0.001), while beyond this (Q2-Q4) the tendency is significantly negative (F_(1, 795)_ = 5.97, p = 0.015). The ‘unlikely’ zone in grey depicts the area of high saccade magnitudes following very short fixations, which should be highly unlikely given the low threshold drive must accumulate needed to break fixation in these instances.

### The oscillator phase influences fixation duration

Having explored saccade amplitude, we next asked whether the model also accounts for fixation duration. In our model, fixations during scanning are periods of fully inhibited locomotion. A fixation ends when the angular drive (generated by the oscillator) crosses a threshold of angular drive (left or right), breaking the inhibition and triggering a saccade (Figure 4A–C). Since the angular drive (L - R activity of the LAL) fluctuates cyclically with the oscillator phase, the time required to reach this threshold naturally varies.

Notably, the models’ fixations can be especially prolonged during reversal phases; when the oscillator transitions from left to right, or vice versa. In extreme cases, the angular drive initially builds towards the direction of the previous saccade (e.g., left), but before crossing the threshold, the oscillator reverses phase. The drive then shifts in the opposite direction (e.g., right) and must accumulate again in that new direction before triggering a saccade (Figure 4B, yellow period of second reversal). As a result, reversal fixations have a tendency to last longer than other fixations (Figure 4A,D). Remarkably, ant behaviour aligns with the model’s prediction (Figure 4E). In our linear mixed-effects model (LME), reversal fixation status was the only significant predictor of fixation duration (*F*_(1,1894)_ = 76.01; *p* < 0.001).

The model also predicts a subtler pattern: fixation duration gradually decreases with distance from a reversal fixation (Figure 4D). This occurs because angular drive is on average strongest mid-cycle (i.e., in the middle of a left or right phase) and weakest near phase transitions. When we replaced the binary “reversal/non-reversal” variable with this sequential phase information (Figure 4A) in our statistical model, the effect on fixation duration remained significant, but only as an interaction with the upcoming saccade’s direction relative to the goal (*F*_(1,1894)_ = 4.65; *p* = 0.031). Specifically, because the CX upregulates the oscillator on the side towards the goal, the angular drive raises quicker and fixation duration tends to be shorter for saccades in that direction.

Together, these results support the idea that fixation duration arises from a threshold-crossing mechanism and signal governed by the oscillator’s phase and its interaction with CX input. Despite outward stillness during fixation, oscillator and CX are continuously active and shaping scan dynamics.

### Influence of neural noise and thresholds on fixation duration

Our model predicts a general negative correlation between fixation duration and the amplitude of the subsequent saccade (Figure 4F, red). This arises because a short fixation (that is, a rapid threshold crossing) implies a strong angular drive, and stronger angular drive typically results in a larger saccade.

However, we were initially surprised to observe in our model that this negative correlation breaks down for the shortest fixations (Q1): in this regime the agents exhibit a positive correlation instead (Figure 4F, blue). This pattern is an indirect consequence of neural noise, which is systematically introduced in the model. Specifically, noise in the inhibitory neuron freezing movement causes variability in the fixation-breaking threshold (illustrated as a grey band in Figure 4B). When this threshold is stochastically lowered due to noise, it can be reached more quickly, producing very short fixations if the current angular drive is strong. However, because the threshold is lower, the accumulated angular drive at threshold crossing is necessarily reduced, limiting the magnitude of the resulting saccade. This mechanism imposes an upper bound on the accumulated drive, and therefore on saccade amplitude, for the shortest fixations, creating a hard limit in the correlation plot (grey region, Figure 4F). The outcome is a positive correlation between fixation duration and saccade amplitude for the shortest fixations (Figure 4F, blue).

While this effect was not initially anticipated, its predictions are supported by the behavioural data. Ants’ fixation duration is positively correlated with saccade amplitude for the shortest fixations (Figure 4G, blue, Quartile 1: *F*_(1, 281)_ = 11.00, *p* = 0.001; *R²* = 0.038; *t*_(281)_ = 3.32, *p* = 0.001), and negatively correlated for longer fixations (Figure 4G, pink, Quartiles 2-4: *F*_(1, 795)_ = 5.97, *p* = 0.015; *R²* = 0.007; *t*_(795)_ = –2.44, *p* = 0.015).

The emergence of this complex pattern from the model, despite not being explicitly designed to capture it, supports the hypothesis that fixations are governed by an internal accumulation of excitatory activity (*angular drive*), which is released upon reaching a threshold. This mechanism is consistent with integration-to-bound neural models that have been proposed for self-initiated actions in other species (e.g. Murakami *et al*., 2014).

### Random Timing of Scan Starts and Stops

Previous work on scanning behaviour in ants (*Deeti et al., 2*023a) found that the number of saccades within each scanning bout follows a Poisson-like distribution (Figure 5B). From this, they inferred that the initiation and termination of scans are governed by a random-rate process.

**Figure 5.**
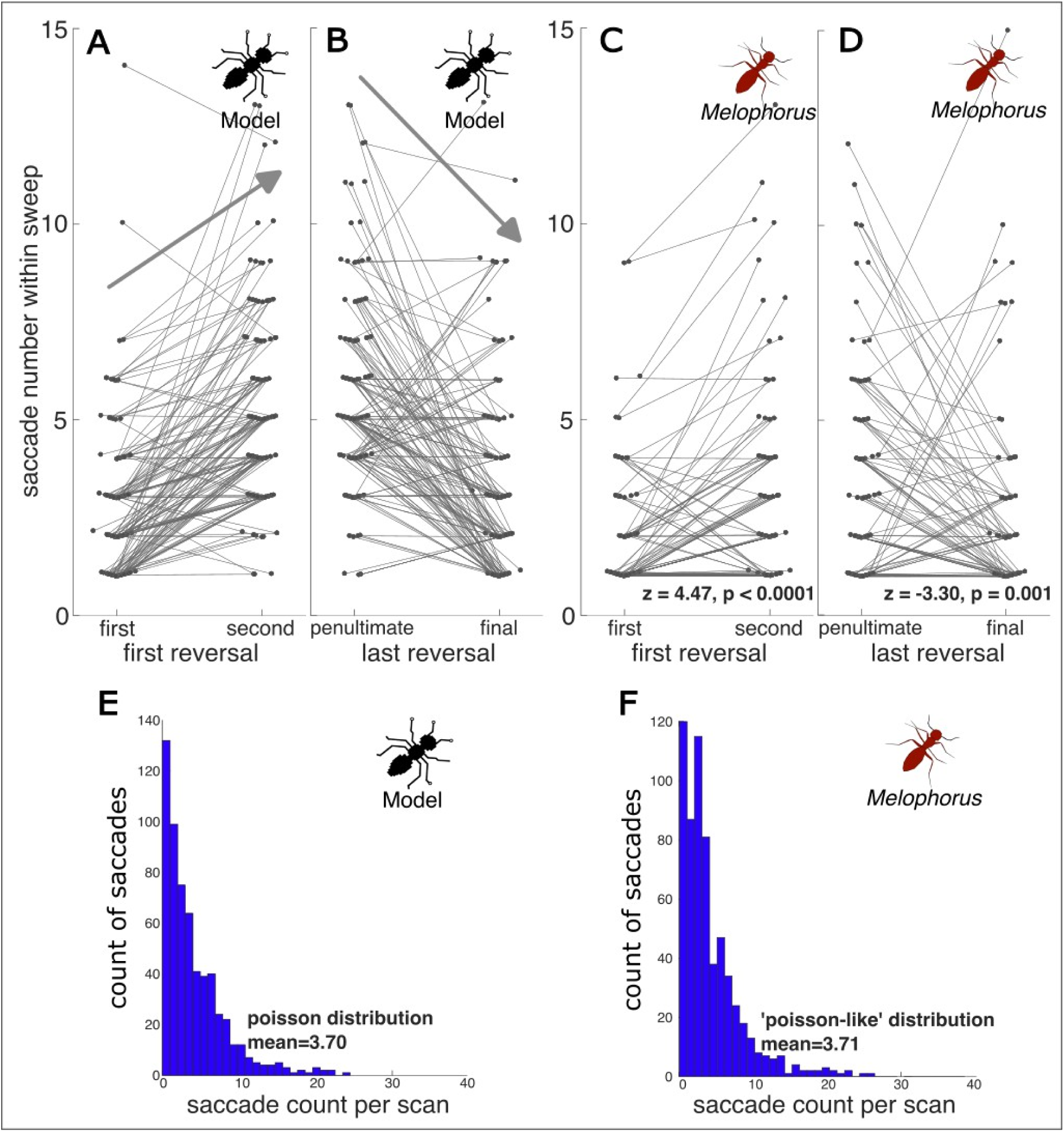
Scans start and end irrespective of oscillator state and saccade number distributions. In the modelled agent, (**A**) the number of saccades within a scanning sweep (before first reversal) was compared with the post-reversal sweep number showing a general upward trend. (**B**) the number of saccades within the penultimate sweep compared to the final sweep of the scanning bout, showing a downward trend. In real ants, (**C**) comparison of the number of saccades in the first and second sweep and (**D**) the penultimate and final scanning sweeps. Both model and real ant sweep comparisons only contain scans which contained two or more reversals, in order to compare the starting/stopping sweeps with a full, within reversals sweep (second/penultimate). Statistical comparisons were made using Wilcoxon tests. The distribution of saccade counts in each scanning bout within (**E**) the model, showing a Poisson distribution and in (**F**) real ants, showing a ‘Poisson-like’ distribution.

In our model, this stochasticity is implemented by a probabilistic trigger: scan initiation is triggered at a random time, and at each moment, scanning can end with a probability proportional to (*1 / mean observed scan duration in ants*), producing the desired Poisson distribution of number of saccades (Figure 5A). Because of this, scans can begin or end at arbitrary points in the oscillatory cycle. This leads to a specific prediction: in scans containing multiple reversals (direction changes in turning), the first and last sweeps, defined here as a bout of sequential saccades in the *same* direction, should often be truncated compared to “full” sweeps (e.g., the second or penultimate sweeps), which start and end exactly at reversals. In other words, sweeps bounded by two reversals should reflect a full (one-sided) oscillatory cycle, while sweeps bounded by the beginning or the end of a scan may not, due to the stochastic start or end of scans. Therefore the latter should tend to be shorter, and thus contain less saccades, than the former.

To test this, we excluded scans with one or zero reversals, focusing on bouts where we could compare truncated sweeps (first and last) with adjacent full sweeps (second and penultimate). The model predicts, and the ant data confirm, that truncated sweeps contain fewer saccades than their adjacent full sweeps (Figures 5A–D). Specifically, the first sweep is significantly shorter than the second (Figure 5C, *z* = 4.47; *p* < 0.0001), and the last sweep is significantly shorter than the penultimate (Figure 5D, *z* = −3.30; *p* = 0.001).

These results thus support the stochasticity of scan initiation and termination. The inhibition of forward motion seems to occur independently of the oscillatory phase. Also, it suggests that within scans, saccade direction follows the underlying oscillatory cycle, rather than being purely random.

### CX strength links navigational uncertainty to scan structure

We next examined how navigational uncertainty, which can be modelled as the strength of the CX’s steering signal, influences scan structure. In the model, a heavily weighted CX signal, representing high certainty in the goal direction, produces greater corrective steering, thus constraining the agent heading towards the goal. As this signal weakens, so does its corrective influence, allowing for larger deviations away from the goal direction. Thus the model predicts an inverse relationship between CX strength (i.e., navigational certainty) and this divergence, which we quantified by measuring the rotation away from the goal covered by the scanning ant before performing a first reversal (Figure 6A, arrow). Ant behavioural data echoes this pattern: experienced ants, which are expected to possess more reliable goal-direction information, showed smaller divergences from the goal on their first reversal than inexperienced ants (Figure 6B; *F*_(3, 274)_ = 3.20, *p* = 0.0239).

**Figure 6.**
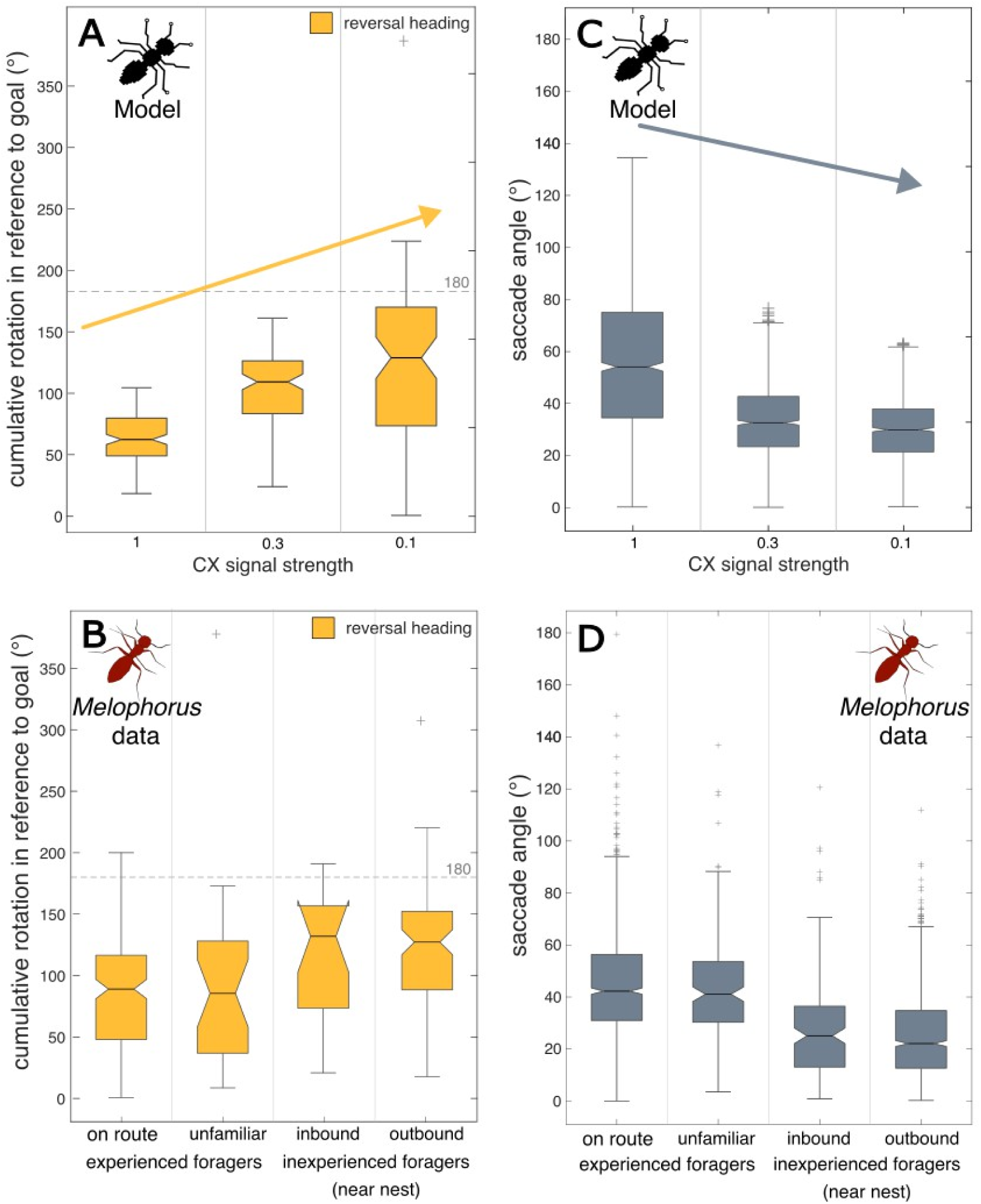
The strength of the CX’s corrective turn signal, and its inverse relationship with navigational uncertainty. The model predicts an inverse relationship between CX signal strength and navigational uncertainty. Signal strength is predicted to be high when navigational uncertainty is low, such as in experienced foragers, and low when uncertainty is high, such as during route formation. (A) Model-predicted first reversal direction (relative to goal direction) as a function of CX signal strength. (B) First reversal direction in real ant data, comparing experienced and inexperienced foragers. (C) Model-predicted saccade angle as a function of CX signal strength. (D) Saccade angle amplitude in real ant data, comparing experienced and inexperienced foragers. For all box and whisker plots, the box spans the inter quartile range while the horizontal line indicates the mean. Whiskers extend to the IQR X 1.5 while outliers beyond this range are shown as ‘+’ symbols. The indentation of the bar around the mean indicates the 95% confidence interval.

The model also predicts a positive association between CX strength and the agent’s saccade angle (Figure 6C, arrow). While initially counterintuitive, as a strongly weighted CX signal constrains angular divergence away from the goal, strong CX steering also produces much larger corrective saccades when it aligns with the oscillator phase (as shown in Figure 3C). This pattern aligns well with our real ant testing conditions, with significantly larger saccades in experienced ants vs. inexperienced ants (Figure 6D; *F*_(1, 2137)_ = 587.6, *p* < 0.001).

These results support a role for the CX’s corrective steering input in modulating the structure of scanning, tuning how tightly ants orient to their goal as well as how wide their saccades are during scanning, especially when in phase with the oscillator.

### Rare ‘Full Loop’ Scans reveal CX–Oscillator Interactions

Despite its simplicity, our model’s closed-loop dynamics generate a surprising diversity of scan forms. One striking example is the emergence of rare ‘Full Loop’ scans (Figure 7A,B; Video 4,5), in which the agent completes a long series of saccades in the same direction, producing a full loop before resuming normal movement.

**Figure 7.**
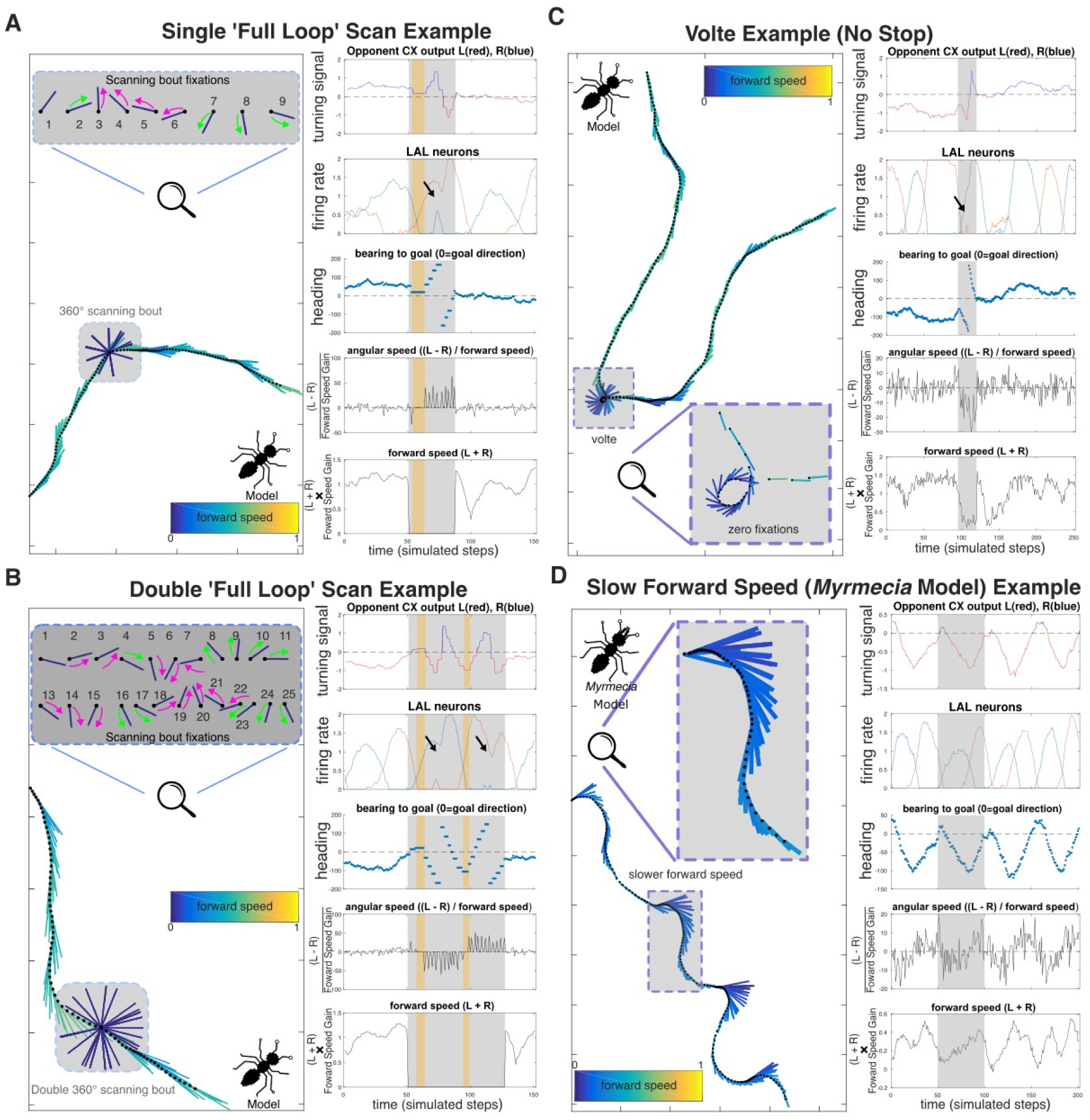
Diverse behaviours emerge from interactions between the central complex’s (CX) steering signal and the modulation of the agent’s forward speed. (**A**) A single ‘Full Loop’ scan (Video 5), the agent terminates forward speed and exhibits a scanning bout with both fixations and saccades. Here, the CX’s corrective steering excites the oscillator and reverses its phase (black arrow), resulting in the agent continuing the loop rotation rather than reversing. These full loop behaviours are rare within scanning (most scans reverse) but are observed in ants (Video 1-3). (**B**) Shows a double ‘Full Loop’ scan example where the agent performs two full loop rotations in a scan. As in panel A, (Simulations - Video 6). Here, the oscillator’s phase is reversed by the CX twice (black arrows). (**C**) A ‘volte’ is a full rotation performed without stopping forward movement, unlike the full loop in A and B.” (Simulation - Video 7). A common behaviour in *Cataglyphis* desert ants (Fleischmann et al., 2017). This behaviour arises in the agent when forward speed is low but not stopped, boosting angular speed and allowing the CX steering output to reverse oscillator phase without fixations. (**D**) An example of a ‘*Myrmecia*-like’ path in the agent, characterized by moderate forward speed and larger lateral oscillations, reminiscent of real world Myrmecia ants (Clément, Schwarz and Wystrach, 2023; e.g. Video - 4). Here, moderate forward speed allows for larger alternating turns and lateral displacement.

In the model, these occur when a strong central complex (CX) corrective steering signal coincides with a critical point in the oscillator’s cycle, effectively shifting the oscillator’s phase by producing a rebound of activity before the current cycle ends (Figure 7A,B, black arrows). Instead of reversing direction at the usual reversal points, the agent’s saccade angle decreases (as usual) but then increases again in the same turning direction, producing a complete rotation.

Such Full Loop scans are not unique to the model; they also occur, albeit rarely, in real ants, and have been reported in multiple species (e.g., *M. bagoti* Figure 1A, Video 1; *Cataglyphis cursor,* Video 2; (Fleischmann et al., 2017; Freas et al., 2019; Freas and Cheng, 2025; Zeil and Fleischmann, 2019).

In the model, full-loop scans arise when CX steering input alters oscillator dynamics sufficiently to prevent the expected reversal. If, by contrast, the CX and oscillator acted purely in parallel (with their outputs summed independently), the oscillator would produce a reversal in angular gain as usual, conflicting with the CX output and preventing a fast completion of the loop. Consistent with the model, real ants show re-acceleration in the second half of the ‘Full Loop’ scan (real ants: Figure 1A, fixations 12–16; Videos 1–3; model: Figure 7A,B), supporting the idea that CX output is indeed branched upstream of the oscillators, and can shift its phase.

These events are too rare in our current dataset for statistical analyses, and our simple model is not designed for quantitative comparisons. Nonetheless, such rare behaviours may offer valuable insights into the coupling between oscillatory motor control and upstream mechanisms, and could be a promising focus for future studies.

### Graded Forward Speed Control Generates Voltes and Oscillatory Patterns

In our initial implementation, forward speed inhibition was binary: the agent ran or stopped completely, as if the central pattern generator controlling leg movement were switched off at a random time. Insects, however, show more nuanced control. For example, desert ants can slow down as their path integration vector decreases (Buehlmann et al., 2018), or accelerate when on fully familiar routes (Clément et al., 2023; Haalck et al., 2023).

We therefore modified the model to allow forward speed inhibition to work along a continuum, with low inhibition merely slowing down the insect rather than stopping it. Because we assumed that forward speed biomechanically impedes angular speed, a sudden but partial reduction in forward speed produces sharper turns; and in some cases complete loops (Figure 7C) resembling so-called ‘voltes’ that have been described in the ant literature (Fleischmann et al., 2017; Zeil and Fleischmann, 2019).

Furthermore, it is known that the oscillator phase itself modulates forward speed, which can be modelled simply as the sum of the oscillator’s left and right activities (Clément et al., 2023). This coupling accelerates the agent when facing its travel direction and slows it when oriented sideways, producing efficient trajectories to cover ground while looking on the sides. The expression of this control varies among species and is particularly strong in *Myrmecia croslandi* (Clément, Schwarz and Wystrach, 2023; Video 4; see also Video 3 for *M. nigriceps*). In the model, operating continuously at lower forward speed releases the constraint on angular speed and naturally produces the large, regular oscillations seen in these ants (Figure 7D) (See Supplemental Figure 1 for example paths).

Interestingly, in such slower agents, increasing the strength of the CX’s influence on the oscillator produces not only more goal-oriented, faster trajectories, but also increases oscillation frequencies while decreasing their regularity (Supplemental Figure 1A,B). These four covarying properties mirror the behaviour of *M. croslandi* ants on familiar routes compared to unfamiliar terrain (Clément et al., 2023), consistent with the idea that visual familiarity generates strong CX goal-heading signals (Wystrach, 2023).

Finally, in strongly oscillating situations, full loops can also arise without apparent external inhibitory signal. Instead, they can emerge naturally from the continuous oscillatory control on forward speed, which slows the animal when facing away from its goal and thus enhances the rotation at the end of a sweep (e.g., *Myrmecia nigriceps*, Video 3). This suggests that the forward–angular speed coupling is a general property of the locomotor system, not solely tied to an external “stop” phase, though whether it is purely biomechanical or neurally reinforced remains to be tested.

Overall, introducing graded forward speed control shows that the same CX–oscillator interactions, originally modelled for scanning, can also explain patterns observed in continuous navigation (periods of goal-directed forward movement), linking route familiarity, speed modulation, scanning behaviours, CX and oscillatory turning dynamics.

## General Discussion

This study shows that diverse movement behaviours in ants, including scanning, pirouettes, voltes, and goal-directed runs, can all emerge from a single, conserved neural architecture. By combining biologically grounded models of central complex steering, intrinsic lateral accessory lobe oscillators, and assuming a physical constraint of forward speed onto angular speed, our models show that modulation of forward speed can capture fine-grained scanning dynamics that match high-speed behavioural data. Importantly, these behaviours do not require a dedicated scanning module; instead, scans,together with a range of observed intermediate behaviours, arise naturally from continuous control mechanisms already implicated in insect navigation. More broadly, these findings suggest that scan-like behaviours need not be interpreted as specialised routines for acquiring nest-aligned views.

### CX and Oscillator Interactions

In our model, the central complex (CX) outputs modulate the oscillator circuit in the lateral accessory lobes (LAL), producing goal-oriented oscillatory paths (Figure 2D; Figure 7A-D, for all paths goal direction to the right). While the CX–LAL pathways have been modelled (Adden et al., 2022) in the context of plume-tracking in moths, their potential implication in ant navigation remains unexplored. The scanning dynamics observed here constrain possible architectures and offer insights into how the CX output may shape motor control via the LAL.

#### CX steering gain increases with angular deviation

Our data show that corrective saccade amplitude increases with the agent’s angular deviation from the goal (Figure 3) which is consistent with behavioural data in ants and flies during continuous navigation (Lent et al., 2010; Westeinde et al., 2024) and suggests that the CX output signal increase with angular deviation. In *Drosophila*, this signature has been proposed to involve PFL2 neurons, which respond maximally when the fly faces away from the goal, and modulate steering gain by converging with PFL3 neurons (which drive left or right turns) onto downstream descending neurons (Westeinde et al., 2024). Here, for the sake of simplicity, we only modelled the two PFL3 populations (PFL-L and PFL-R), tuned to both ±135° from the goal (i.e., ±45° from the anti-goal), which also mimics CX steering cells observed in monarch butterflies (see Fig.4G of Beetz, Kraus and el Jundi, 2023), and result in similar steering dynamics as observed in *Drosophila* (compare Figure 2A and Westeinde et al., 2024). Whatever the exact implementation, the preserved relation between amplitude and angle away from the goal during scan (Figure 3) supports the idea of a conserved CX output function that encodes angular deviation from the goal with increasing strength, and is at play during both scanning and continuous navigation.

#### CX directly modulates the LAL oscillator

A key question is whether the CX directly modulates the LAL oscillator (serial control), or whether both act independently in parallel.

In moths, descending neurons in the LAL exhibit characteristic ‘flip-flop’ activity patterns that correlate with zigzagging maneuvers (Olberg, 1983; Kanzaki and Ikeda, 1994). Computational models suggest that having these LAL neurons modulated by the CX output can explain aspects of the moths’ plume-tracking behaviour (Adden et al., 2022). In ants, behavioural studies show that strong directional drives elicited by the path integrator or visual familiarity do not only gain behavioural weights and sharpen directional accuracy (Wehner et al., 2016; Wystrach et al. 2015, Legge et al. 2014) but also increase the ants’ oscillation frequency (Haalck et al., 2023, Clément et al., 2023). Assuming that the path integrator and visual familiarity modulate goal signals in the CX, as modelled here and elsewhere (Wystrach et al., 2020b, Stone et al., 2017) and that the intrinsic oscillator is in the LAL (Clément et al., 2023, Steinbeck et al., 2020), this frequency increase suggests that CX output modulates the intrinsic oscillatory activity of the LAL. Similar increases in ants’ oscillation frequency also occur in response to optic-flow prediction errors, although these signals may bypass the CX (Dauzere-Peres and Wystrach, 2024).

Our findings offer a qualitatively different form of evidence for a direct modulation of the LAL oscillator by the CX. The rare ‘full loop’ scans - 360° turns without reversal - only emerge in the model because the CX can shift the oscillator’s phase mid-cycle (Figure 3A,B). This requires tight coupling between CX output and LAL dynamics, and would not occur if both acted independently. Together, present and past results provide support for a direct, serial modulation of the LAL oscillator by the CX in ants.

#### An opponent process in the CX output

In our model, the output of the CX to the LAL (via PFL neurons) provides both ipsilateral excitatory connections and a contralateral inhibitory connection (Figure 2A). In insects, the latter may be indirect, via the stimulation of inhibitory neurons such as the protocerebral bilateral neurons in the LAL (Kanzaki et al., 2004; Mishima and Kanzaki, 1999). Functionally, this contralateral inhibition ensures that only one side of the LAL is activated by the CX at a time, avoiding simultaneous bilateral activation, which in the model would produce inappropriate bursts of forward motion, particularly when the agent is misoriented.

More generally, this opponent organisation between the left and right hemisphere converts CX directional information into a normalised steering signal that reflects angular deviation. Similar opponent processes have been proposed for mushroom body output neurons encoding approach vs. avoidance, or left vs. right scene familiarity signals (Le Möel and Wystrach, 2020; Murray et al., 2019; Wystrach, 2023), suggesting that lateralised, opponent coding is a general principle in insect navigational circuits for ensuring stable and interpretable motor outputs.

## A Simple Control Principle: Forward Speed Gates Exploration

A key insight from this work is the importance of the assumed constraint of forward speed on the expression of body rotation. This effectively enables a one-dimensional control to adjust the expression of active sampling along the tradeoff between exploitation and exploration. At one extreme, forward speed is fully inhibited, halting leg CPG output, and making angular drive maximally expressed, producing what we term scans. At the other, high forward speed suppresses angular turns, yielding straight, fast goal-directed trajectories. By simply adjusting this one parameter, the model generates a spectrum of behaviours, from sweeping oscillations to full rotational scans, mirroring inter- and intra-species variation observed in real ants (Figure 8). This spectrum of apparently distinct behaviours does not reflect different mechanisms, but the same distributed control process. The diversity arises from the dynamic interactions between the CX, the oscillator, and thoracic CPGs, and, crucially, the mechanical constraint of forward speed on rotation.

**Figure 8.**
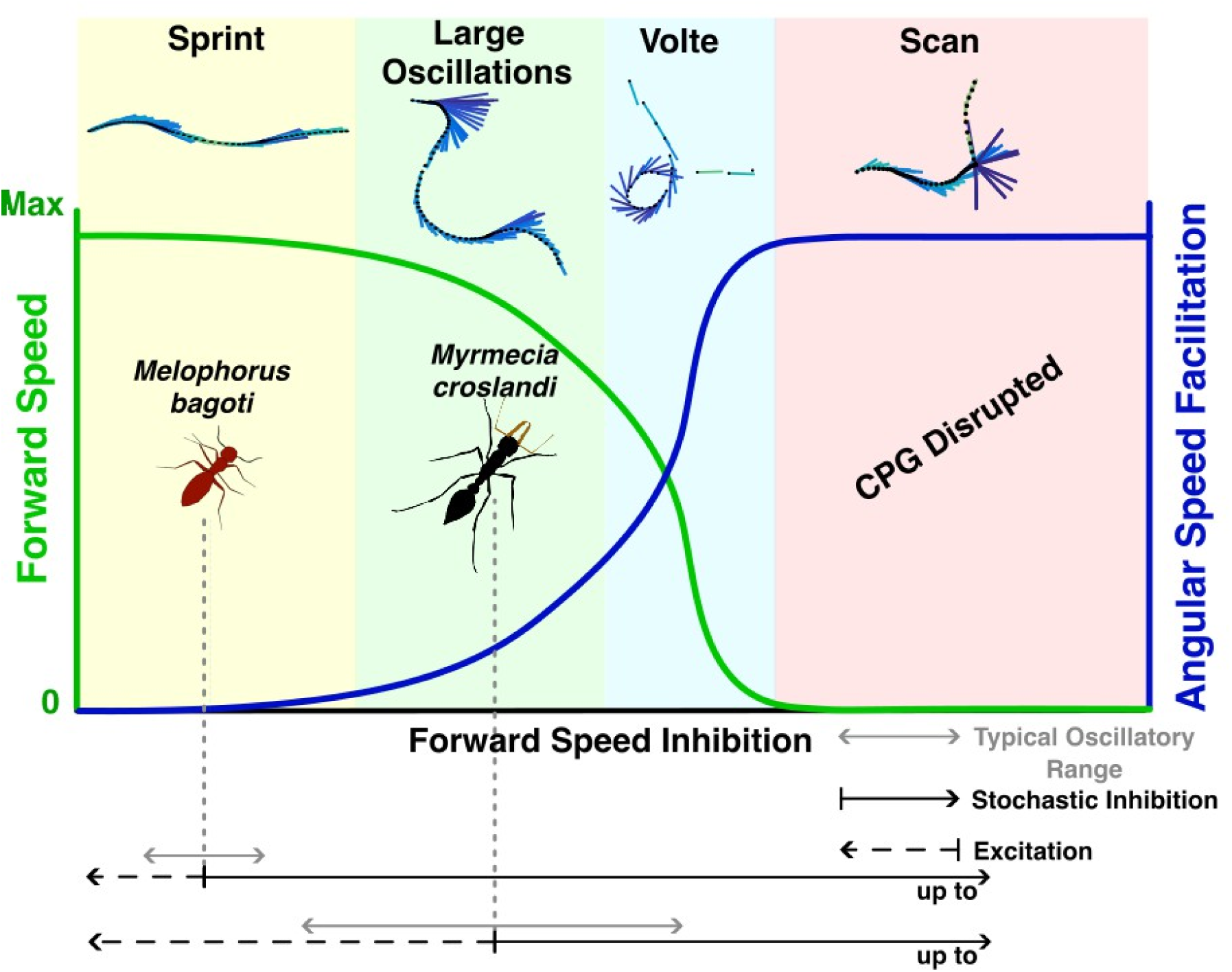
Summary of the behavioural spectrum produced by the modelled agent as a function of forward speed inhibition facilitating angular speed. High forward speed results in an agent that ‘sprints’, with its straight paths resembling desert ants (*Melophorus bagoti and Cataglyphis*). Decreasing forward speed progressively increases angular speed facilitation, leading to the large oscillations of Myrmecia under moderate forward speed (Clément, Schwarz and Wystrach, 2023; Video 4), ‘voltes’ under low forward speed, and ultimately scanning behaviours when forward speed reaches zero. Here, angular facilitation plateaus, the central pattern generator (CPG) is disrupted and the agent scans, with the choreography of scanning being dictated by interactions between CX signal strength and the oscillator, as observed in ants (Real Ants - Video 1-3; Simulations - Video 5,6)

### A continuum of movement modes across context and species

The continuum also helps explain interspecific variation. Desert ants (*Melophorus bagoti,* multiple *Cataglyphis species, Occymyrmex robustior*), adapted to thermophilic foraging, favour high forward speed, which thus limit oscillation amplitudes, and rely on brief, stochastic stops for visual sampling (Clément et al., 2023; Deeti and Cheng, 2025; Muser et al., 2005); figure 8). *Myrmecia* ants, by contrast, forage at lower temperatures and can afford to operate at slower speeds, allowing large, regular oscillations to coexist with forward progression (Clément et al., 2023); Figure 8). Our model can shift seamlessly from one style to the other by simply adjusting forward speed.

An individual can also shift along the continuum to adapt to the current context. For instance, experienced *Cataglyphis velox* running along their familiar route, where navigational certainty (implemented as a strong goal heading in the CX, Figure 6), favour exploitation by constraining angular deviation, increasing forward speed and thus reducing the occurrence of scans. Similarly, *Myrmecia* can stop and scan under high uncertainty (Freas et al., 2018; Islam et al., 2020) or accelerate and straighten to escape adverse situations (Clément et al., 2023; Deeti et al., 2023b).

The emergence of such different-looking trajectories suggests that the modulation of forward speed may represent an ancestral and fundamental control strategy in ants. This simple control is used at both individual decision and evolutionary time scales: to adjust individuals’ behaviour to the current context, as well as ant species to their ecology.

### An ancestral design? Striking parallels with crawling Drosophila larvae

This forward–angular coupling is not unique to ants. Similar mechanisms appear across taxa (even in bacterial run–tumble cycles), highlighting a shared control strategy (Cheng, 2024). Importantly, while these similarities do not imply homologous neural implementations, they do suggest that comparable behavioural motifs can emerge from related control principles across taxa. In *Drosophila* larvae, angular reorientation also arises from a continuous body-bend oscillator. The expression of body bending is constrained by forward crawling peristalsis (Wystrach et al., 2016), which can be transiently suppressed by descending neurons (e.g., PDM-DN), producing a stop and unmasking the oscillator expression by enabling strong body rotation (Tastekin et al., 2018; Berni et al., 2012; Pulver et al., 2015), akin to ant scans.

Fly larvae’s stops also appear stochastic, and their frequency can be modulated by negative sensory evidence such as going down attractive odour gradient (Berni et al., 2012; Gomez-Marin and Louis, 2014), just as scanning in ants is more likely under high navigational uncertainty (Deeti et al., 2023a; Freas et al., 2022, 2018; Freas and Cheng, 2025; Schwarz et al., 2020a; Wystrach et al., 2020a, 2014).

Finally, in both ants (Buehlmann et al., 2018; Haalck et al., 2023) and larvae (Gomez-Marin and Louis, 2014; Luo et al., 2010), forward speed control is not binary, but graded, modulated smoothly by sensory evidence.

Although not present as neuropils in fly larvae, the CX and LAL are conserved structures, ancestral to arthropods (Kanzaki, 2005; Kanzaki and Mishima, 1996; Pfeiffer and Homberg, 2014; Heinze, 2024). Whether the descending neurons that inhibit forward movements in *Drosophila* (Rayshubskiy et al., 2025; Tastekin et al., 2018) are similarly conserved across arthropods, or represent convergent evolutions in some taxa, remains to be seen.

In adult *Drosophila* and other insects, multiple descending neurons modulating forward speed have been identified (Büschges and Ache, 2025). Some (BPNs) promote acceleration (Bidaye et al., 2020), whereas others trigger strong speed reductions, or halting responses (Sapkal et al., 2024). Furthermore, left-right firing rate differences between descending neurons (DNa-1/DNa02) can predict rotational velocity driving pivoting behaviour (Büschges and Ache, 2025; Rayshubskiy et al., 2025; Yang et al., 2024); providing a circuit-level analogue to our modelled angular speed modulation.

### Scanning in flying hymenopterans

Flying hymenopterans such as bees and wasps share much of their navigational toolkit with ants (Collett et al., 2013; Collett and Hempel de Ibarra, 2023; Zeil, 2012). As in ants, their scene recognition and path integration systems are thought to generate a goal heading within the CX (Stone et al., 2017; Honkanen et al., 2019) and they also exhibit regular oscillations and scanning behaviours, particularly during learning flights or when approaching a goal (Philippides et al., 2013; Collett et al., 2013, 2023; Zeil et al., 1996). However, these behaviours are expressed differently from those of ants, notably through sideways motion, arcs, and loops, rather than by simply stopping and rotating on the spot.

The bodily constraints assumed in the present model are intended to reflect those of a typical walking ant, thus the agent cannot move sideways. Ants are capable of sideways or backward motion, but apparently only when no alternative is available, such as when dragging a heavy item (Schwarz et al., 2017). It would be interesting to determine whether scanning properties such as those observed in bees and wasps could emerge from the model proposed here under different physical constraints adapted to flight, while keeping the same neural mechanisms. Novel behaviours may indeed evolve through changes in the body rather than in the brain (Dudley, 2000).

### Model limitations and future directions

While our model successfully captures fine-scale scan dynamics, several limitations remain. First, we assumed a single, static goal direction represented in the fan-shaped body of the CX. While this suffices to simulate realistic scanning of ants navigating towards a known goal, real insects often juggle multiple directional cues, from path integration, visual landmarks, wind, and olfaction, and may maintain simultaneous goal representations. These goals may be employed sequentially or weighted adaptively within the CX (Freas et al., 2020; Le Moël et al., 2019; Stone et al., 2017; Sun et al., 2020; Wehner et al., 2016; Wystrach et al., 2015). Extending the model to simulate multi-goal conflict or cue integration would allow exploration of how insects handle navigational uncertainty, and how the latter impacts scanning.

In our simulations, the CX goal representation remained fixed in both direction and strength throughout each trial. This simplification allowed us to isolate and compare the effects of different CX strengths on scanning behaviour (Figure 6). However, goal headings in the CX are likely to be updated continuously (including during the scans themselves), for instance via novel inputs from visual recognition in the MB (Goulard et al., 2021; Sun et al., 2020). This would affect saccades direction and duration. Exploring such dynamics lies beyond the scope of the present study but would represent an interesting direction for future work. Notably, our proposed CX-LAL-Body relationship could be implemented downstream of an existing path integration or visual-based model (or both) to form predictions about the occurrence and dynamic of scans along the path, as well as their impact on the emerging trajectories.

The model also does not currently explain the phenomena of nest-directed fixations. These so-called “pirouettes” or “lookbacks”, where longer fixations happen when looking towards the nest have been reported in ants during learning walks or initial route formation (Collett et al., 2023; Collett and Hempel De Ibarra, 2023; Fleischmann et al., 2017; Freas and Cheng, 2018, 2025; Müller and Wehner, 2010; Robert et al., 2018; Stürzl et al., 2016; Zeil and Fleischmann, 2019). While in our data set scans did rarely extend beyond 150° away from the goal (Figure 6), our model predicts that when this goal is weak - such as in entirely naive ants - reversal, smaller saccades and longer fixations will tend towards 180° from the goal, which might coincide with the origin, that is, the nest direction. Alternatively, the nest direction could actually be represented as a goal heading. This could be addressed by incorporating either (a) a dual-goal representation (e.g., one for the feeder and one for the nest) with modulated weighting, or (b) a dynamic switch of goal orientation during the scan, perhaps based on distance from home. This raises several empirical predictions. For example, if ants perform scans while navigating along two-leg outbound routes where the goal is at 90° to the nest’s direction, then fixation patterns during scans may reveal whether multiple goals influence scanning structure. If the CX indeed holds multiple co-active directional representations, scan fixations should systematically cluster around these multiple directions.

Finally, scan termination is currently modeled as a stochastic process, implemented through a random-rate “freeze-release” mechanism to reproduce the observed Poisson-like distribution of scan durations (Deeti et al., 2023a). However, behavioural evidence suggests that scan duration is not entirely random. Scans tend to last longer in contexts of high uncertainty, such as during learning walks or early route formation, and become shorter or absent on well-learned routes (Wystrach et al., 2014). This implies that the probability of termination likely depends on an internal variable, perhaps an accumulating “forward drive” or confidence estimate, that builds over time and releases the motor system from inhibition once a threshold is reached. This mechanism would parallel the angular drive threshold that governs saccade initiation in our current model (Figure 4) and resembles integration-to-bound processes proposed in other species, including mammals (Murakami et al., 2014).

While our model does not include detailed biomechanics of leg coordination, proprioceptive feedback, or thoracic pattern generators (CPGs), it makes testable predictions about how angular and forward drive might interact at the motor level. For instance, the assumption that angular drive and forward drive accumulates continuously, even during motionless fixations predicts the existence of neurons or motor circuits that integrate excitatory input over time and release movement when thresholds are crossed, paralleling the control logic found in *Drosophila* larvae (Rayshubskiy et al., 2025; Tastekin et al., 2018). Whether such a putative integration over time is achieved at the level of descending neurons, or downstream in the thoracic CPG themselves remains to be seen.

### Conclusions

This study demonstrates that the diverse movement patterns of ants, from brief scans to full pirouettes and sweeping oscillations, can arise from a single, conserved neural architecture: the interaction between central complex (CX) steering and an intrinsic oscillator in the lateral accessory lobes (LAL). By introducing minimal, biologically plausible additions (a stochastic inhibition of forward speed, a saccade-initiation threshold, and forward–angular speed coupling), the model reproduces a panel of fine-scale scanning dynamics seen in high-speed recordings of *Melophorus bagoti*, and generalizes across species and behavioural contexts.

A key insight is that forward speed acts as a distributed control dial, gating the expression of angular movements and enabling smooth transitions between goal-driven progression and exploratory sampling. This unifying principle offers a simple yet powerful mechanism for balancing exploitation and exploration, adaptable across ecological niches and timescales.

Together, these directions open exciting opportunities to link brain models of sensorimotor control to peripheral control such as thoracic ganglia, refine our understanding of decision timing, and extend the model to richer multi-goal or multisensory environments.

## Materials and Methods

### Model Overview

We developed a computational model to test whether scanning behaviour dynamics can emerge from the interaction between central complex (CX) steering and an intrinsic lateral accessory lobe (LAL) oscillator. The model simulates single runs where the virtual agent possesses a fixed goal direction in the CX and forward speed will be inhibited at a predetermined time and released stochastically to trigger a single scanning bout. The model was implemented in MATLAB R2016b, and code is openly available at https://github.com/antnavteam/CX_oscillator_scans. The code is extensively commented to allow readers to examine all implementation details, parameter choices, and computational logic.

The model comprises three interacting components: (1) a CX steering circuit that compares current heading with a goal direction to generate corrective signals (adapted from Wystrach et al., 2020b), (2) an intrinsic LAL oscillator that produces rhythmic left-right activity (adapted from Clément et al., 2023), and (3) motor output rules that convert neural activity into angular and forward velocity.

### Central Complex Steering Component

The CX steering architecture follows the computational framework established in previous insect navigation models (Stone et al., 2017; Wystrach et al., 2020b), where current and goal direction are encoded as bumps of activity across rings of 8 neurons, and compared via shifted representations to generate steering commands. The CX encodes heading information as activity patterns across eight sectors with fixed angular references every 45°:

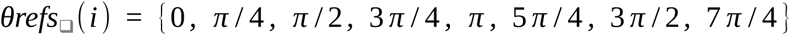

Current heading is represented by a bump of activity across eight EPG (Ellipsoid body-Protocerebral bridge-Gall) neurons, which track the agent heading (TH). The goal direction is represented by a bump of activity across eight goal cells (here termed HDB), which remains fixed at *θ* _ *goal* **=** 0:

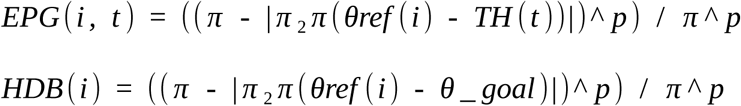

In these equations, TH(t) represents the agent’s current heading, θ_goal is the fixed goal direction (which we set to 0), p (= 3) controls the width of the activity bump, and π₂π() wraps angles to [-π, +π]. Both EPG and HDB produce activity bumps centered on their respective directions, with activity falling off smoothly away from the peak.

Two PFL (Protocerebral bridge-Fan-shaped body-Lateral accessory lobe) populations compare spatially shifted copies of the heading representation with the goal representation. Each PFL population computes the difference between the goal signal and a shifted heading signal at each PB sector. Because neural activity under inhibition is bounded at zero, negative values are prevented and set to zero:

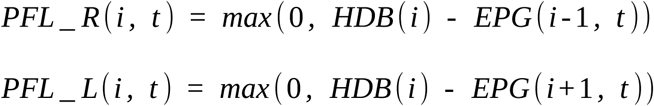

where the indices i-1 and i+1 represent spatial shifts of ±1 sector (±45°). When the goal representation is stronger than the (shifted) heading representation at a given position, the PFL neuron becomes active; otherwise it is silent. This generates asymmetric activity patterns: PFL_R becomes active when the goal lies to the right of the agent’s current heading, and PFL_L when the goal lies to the left of the agent’s heading. The summed activity across all sectors produces the bilateral CX steering output:

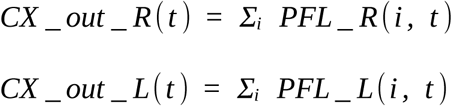

The net difference between these outputs indicate whether the goal lies rather to the right or the left of the current heading. Evidence in butterflies shows that corrective output signals of the CX peaks when the insect is oriented 135° away from its goal (Beetz et al., 2023). We implemented this by simply reversing the CX steering signal (see below): evidence for the goal on the right will steer the agent towards the left (and reciprocally), so that the agent will tend to move in the opposite direction (180° away) from his CX goal representation, which effectively, acts here as an anti-goal. As a result, the comparison of ±1 sector (±45°) shifted representation makes the magnitude of CX_out peaks when current heading is shifted by 45° from the CX goal, that is 135° (180° - 45°) from the agent goal (Figure 3). Note that this is equivalent to a shift of ±3 sectors (±135°) with a non-reversed steering control. As a result, the CX produces a stronger corrective steering signal when the agent is more misoriented, a property consistent with behavioural data in ants and flies (Lent et al., 2010; Westeinde et al., 2024).

To produce unambiguous unilateral steering signals, the two CX outputs go through a cross hemispheric reciprocal inhibition (Figure 3), and then modulates the oscillator via ipsilateral activation: *CX* _ *input* _ *R* for the right oscillator neuron R, and *CX* _ *input* _ *L* for the left oscillator neuron L.

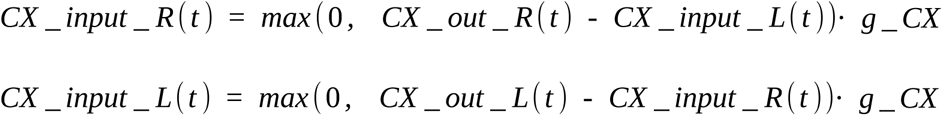

g_CX represents the CX gain parameter, that is, the strength of the CX impact on the oscillator, which remains constant throughout each run, and whose impact on scanning was investigated (Figure 6)

#### Intrinsic LAL Oscillator

Two reciprocally inhibiting neurons (R and L) generate oscillatory activity through mutual inhibition and negative feedback representing synaptic exhaustion (Clément et al., 2023):

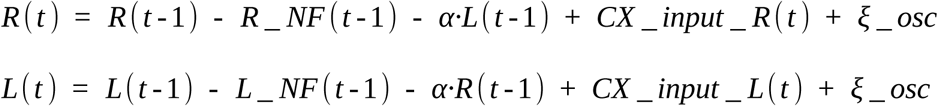

α = 0.1 represents the reciprocal inhibition strength and ξ_osc ∼ N(0, 0.05) is Gaussian noise. The negative feedback (synaptic exhaustion) evolves as:

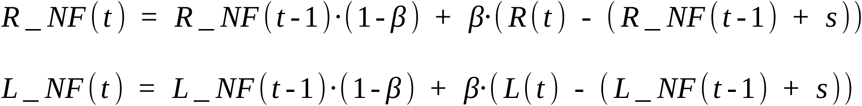

β = 0.01 represents the exhaustion rate and s = 0.5 is the steady-state firing rate. Neural activity is constrained to [0, 2] to prevent runaway firing activity.

#### Motor Output

Angular velocity (ω) and forward velocity (v) are derived from oscillator activity:

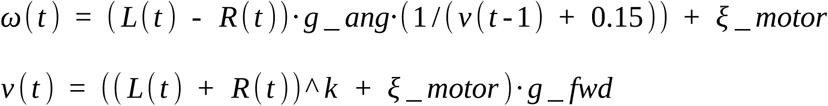

g_ang = 0.03 represents angular motor gain, g_fwd represents forward motor gain, k controls how much forward speed varies with oscillator output, and ξ_motor ∼ N(0, 0.1) is motor noise. The term (1/(v + 0.15)) implements the coupling between forward and angular speed: faster forward movement reduces turning ability. +0.15 prevents an infinite value when v=0.

To simulate species differences, we used two parameter sets:

Desert ants (e.g., *Cataglyphis* and *Melophorus*): k = 0.5, g_fwd = 1.0 (faster, smaller impact of the oscillator on forward speed)

*Myrmecia* ants: k = 1.5, g_fwd = 0.2 (slower, larger impact of the oscillator on forward speed)

The agent’s heading and position update at each timestep:

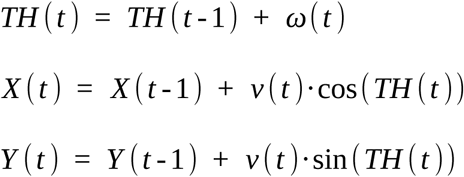

### Scan Generation via Forward Speed Inhibition

Forward speed can be inhibited to trigger scanning behaviour. We implemented two inhibition modes:

**Full inhibition (fwd_speed_inhibition = 0):** Forward velocity is set to zero, halting the locomotor central pattern generator (CPG). During this period, angular drive (*L*(*t*) - *R* (*t*)) accumulates :

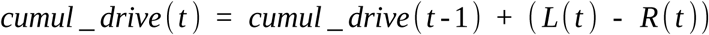

The agent remains in fixation (ω(t) = 0) until accumulated drive exceeds a threshold to break the CPG inhibition θ_CPG = 2.0, then releases a saccade using the usual formula for rotation:

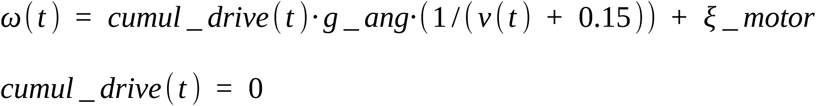

This release resets the cumulated drive to zero, that is below CPG desinhibition threshold, making the agent freeze and the angular drive cumulation resume. This produces the characteristic scan structure of alternating fixations and saccades observed in ants.

**Partial inhibition (fwd_speed_inhibition > 0):** Forward speed is reduced but not halted (e.g., set to 0.2), and angular control proceeds normally without accumulation. This generates broad oscillations or “voltes” rather than discrete scans.

Scan termination is stochastic: at each timestep during inhibition, forward speed resumes with probability p_stop = 0.5. This produces exponentially distributed scan durations matching the Poisson-like distribution observed in *M. bagoti* (Deeti et al., 2023a).

### Model Parameters

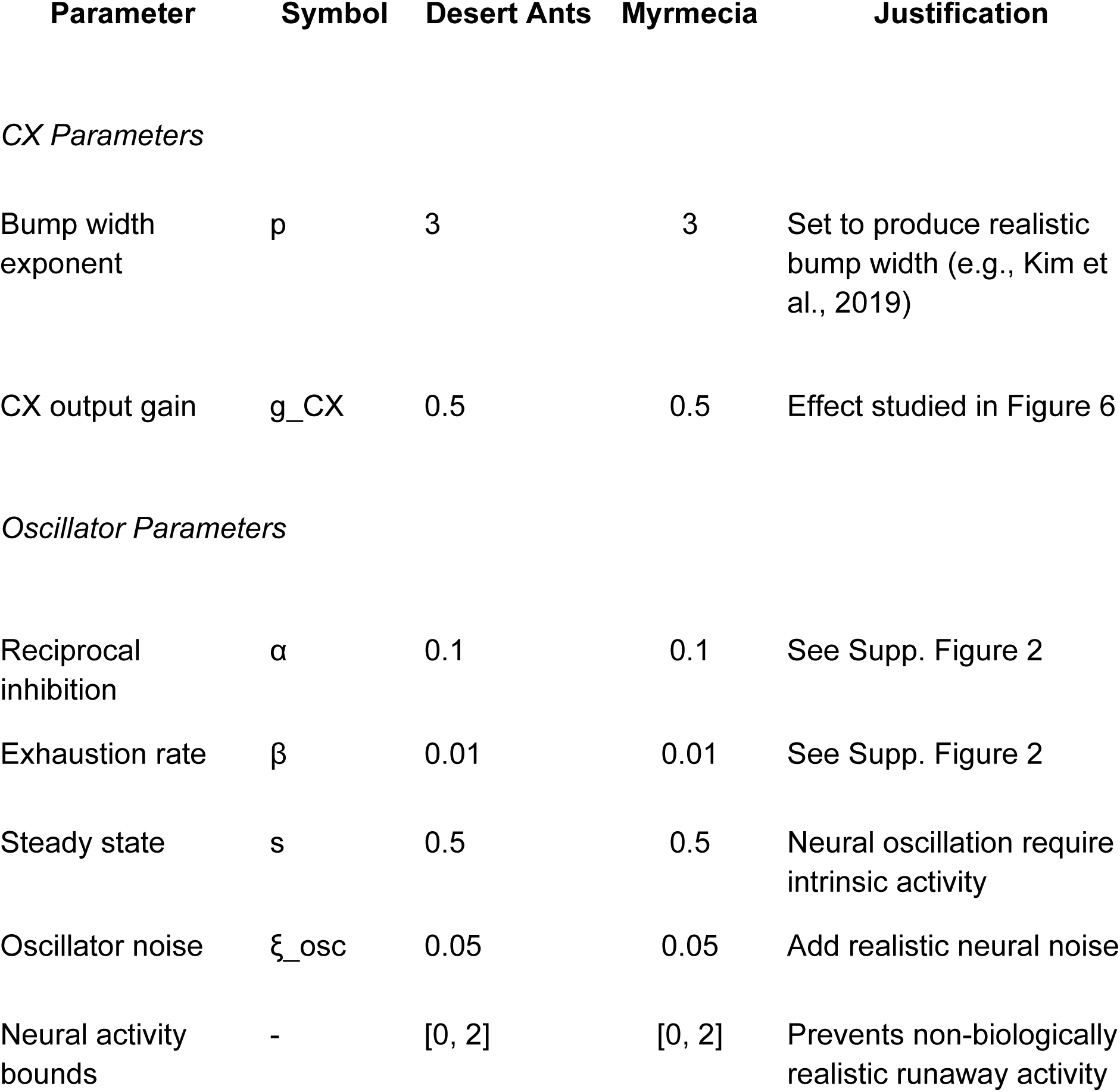
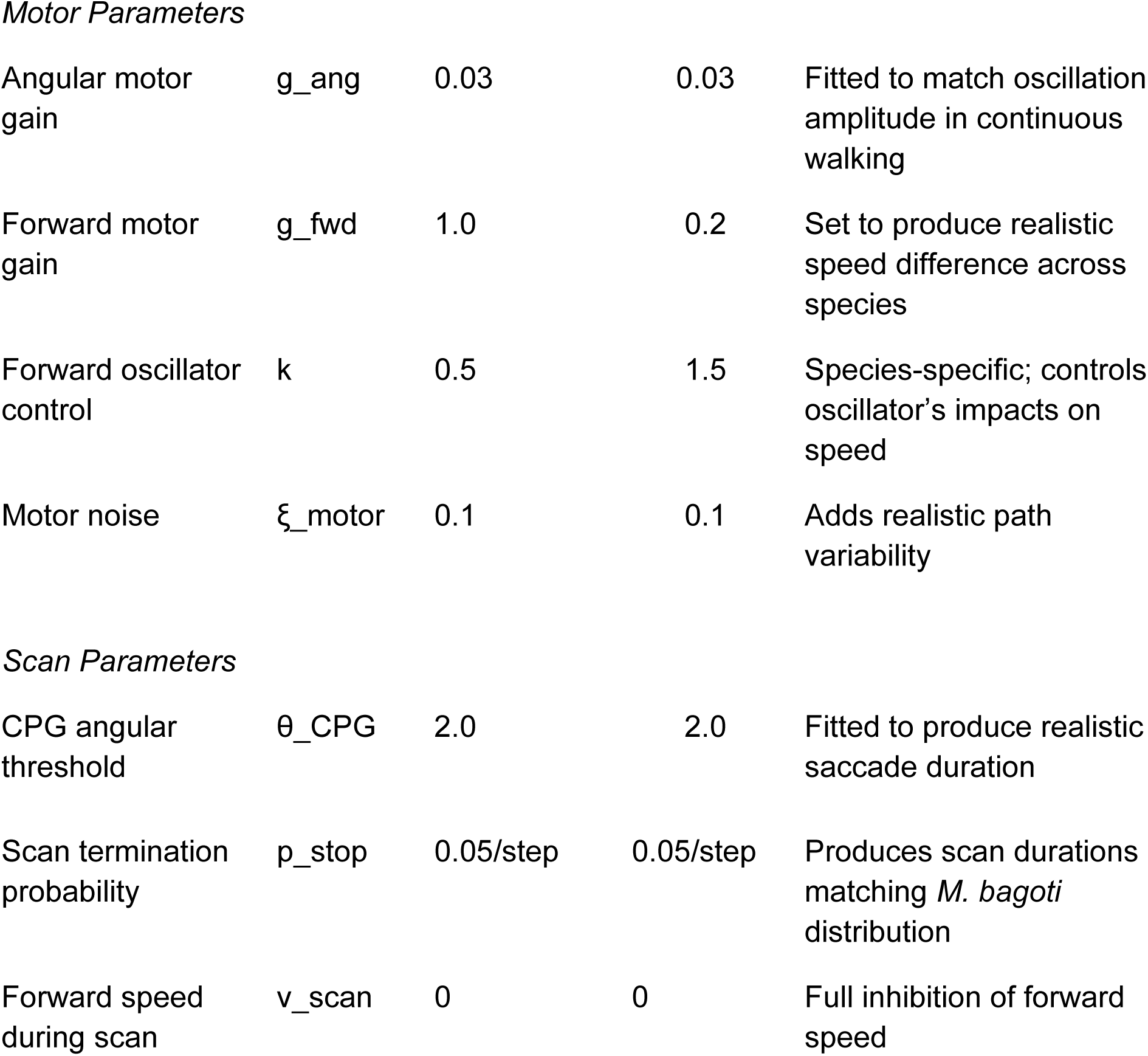

| Parameter | Symbol | Desert Ants | Myrmecia | Justification |
| --- | --- | --- | --- | --- |
| *CX Parameters* |  |  |  |  |
| Bump width exponent | $p$ | 3 | 3 | Set to produce realistic bump width (e.g., Kim et al., 2019) |
| CX output gain | $g_{\text{CX}}$ | 0.5 | 0.5 | Effect studied in Figure 6 |
| *Oscillator Parameters* |  |  |  |  |
| Reciprocal inhibition | $\alpha$ | 0.1 | 0.1 | See Supp. Figure 2 |
| Exhaustion rate | $\beta$ | 0.01 | 0.01 | See Supp. Figure 2 |
| Steady state | $s$ | 0.5 | 0.5 | Neural oscillation require intrinsic activity |
| Oscillator noise | $\xi_{\text{osc}}$ | 0.05 | 0.05 | Add realistic neural noise |
| Neural activity bounds | - | [0, 2] | [0, 2] | Prevents non-biologically realistic runaway activity |
| Motor Parameters |  |  |  |  |
| Angular motor gain | $g_{ang}$ | 0.03 | 0.03 | Fitted to match oscillation amplitude in continuous walking |
| Forward motor gain | $g_{fwd}$ | 1.0 | 0.2 | Set to produce realistic speed difference across species |
| Forward oscillator control | $k$ | 0.5 | 1.5 | Species-specific; controls oscillator's impacts on speed |
| Motor noise | $\xi_{motor}$ | 0.1 | 0.1 | Adds realistic path variability |
| Scan Parameters |  |  |  |  |
| CPG angular threshold | $\theta_{CPG}$ | 2.0 | 2.0 | Fitted to produce realistic saccade duration |
| Scan termination probability | $p_{stop}$ | 0.05/step | 0.05/step | Produces scan durations matching *M. bagoti* distribution |
| Forward speed during scan | $v_{scan}$ | 0 | 0 | Full inhibition of forward speed |

### Simulation Protocol

Each simulation ran for 400 timesteps, representing (given our oscillatory parameter choice, see Supplemental figure 2) around 6 to 10 oscillation cycles (depending on CX strength). This would take ants around 5s to 25s (depending on species, individual, temperature and other conditions), which is sufficient to encompass one realistic scan duration. Initial conditions were: R(0) = 1, L(0) = 0, TH(0) = 0, position (0, 0). Forward speed inhibition (i.e., scanning) began at timestep 200 (±50 timesteps random variation to ensure that scan onset occurs at various moments along the oscillatory cycle). The model generated time series of:

- Position (X, Y) and heading (TH)
- Neural activities (R, L, CX_outputs)
- Velocities (angular ω, forward v)
- During scans: fixation durations, body orientations, saccade amplitudes

### Model Outputs and behavioural Comparisons

From scanning periods (when v = 0), we extracted:

**Fixation metrics:** Duration (time between saccades), body orientation relative to goal direction

**Saccade metrics:** Angular amplitude, direction (toward/away from goal), reversal events (change in turning direction)

**Scan structure:** Total duration, number of fixations per scan, cumulative orientation at first reversal

These were compared qualitatively against empirical data from *M. bagoti* using the same metrics extracted from high-speed videos (see behavioural Experiments section). We did not perform quantitative parameter optimization, as our goal was to assess whether scanning properties naturally emerged from the proposed circuit architecture rather than to fit the model precisely to data.

### Behavioural Experiments

#### Species and site

Natural scans were recorded using the red honeypot ant *Melophorus bagoti,* during the Australian summer from December to February (experienced foragers - *Familiar*/*Unfamiliar* conditions: 2010, see Deeti et al. 2023; inexperienced foragers – *Outbound/Inbound* route formation conditions: 2023/2024 summer season) on a field site (23°45′28.12″S, 133°52′59.77″E) located at the Centre for Appropriate Technology campus, south of Alice Springs, Northern Territory, Australia. *M. bagoti* inhabit a visually cluttered environment, exemplified by large amounts of buffel grass tussocks (*Pennisetum cennchroides*), with scattered eucalyptus trees and bushes.

Desert ants navigate alone using multiple concurrent, primarily visual, strategies (Cheng et al., 2009; Collett and Collett, 2002; Freas and Spetch, 2018, 2023; Wehner, 2009; Wystrach et al., 2012; Zeil, 2023); including path-integration (Cheng et al., 2009; Collett and Collett, 2002; Wehner and Srinivasan, 2003) and learned views around the nest and along foraging routes (Collett, 2010; Freas et al., 2017; Wystrach et al., 2012; Zeil, 2023; Zeil and Fleischmann, 2019).

Two data sets of high speed videos of scanning bouts were used in this project. The first set of high speed scanning bout videos were collected using inexperienced foragers, as individuals formed a straight-line route between a feeder and the nest. In these ants, scanning bouts are known to have a high occurrence near the nest, in both outbound and inbound ants. Videos were collected both at the onset of their *outbound* journey or during the *inbound* trip just before reaching the goal. The second was a group of highly experienced ants which were released either along or near their established foraging route (familiar) or at a distant (unfamiliar) site (previously collected in Deeti et al. 2023).

#### Inexperienced forager procedure

High speed video recordings of inexperienced ants as they scanned while forming their stereotypical route between and nest and feeder conducted on a single *M. bagoti* nest in a closed arena with 10cm high walls enclosing the nest and feeder (Same arena as Freas and Cheng, 2025, with a separate cohort of ants). A stocked feeder was sunk into the ground 7m from the nest. As we were only interested in inexperienced foragers with little knowledge of the area beyond the nest, prior to video recording all foragers that emerged from the nest entrance were marked as experienced using enamel paint (Tamiya™) and these experienced individuals were excluded. On day six, non-painted foragers were allowed to exit the nest entrance and find the feeder. Extending 0.2-1.0m from the nest entrance we spread a thin layer of white sand along the ground. This sand helped the red ants stand out from the red ground during filming and pose estimation analysis. Above this site, we positioned a downward facing Chronos 2.1-HD high-speed camera (KRON Tech), 50cm above the ground (1920×1080pixels, 600fps) with a field of view of 30cm×17cm.

For five days prior to recording, one exposure to the feeder encouraged foragers to leave the nest entrance in the general direction of the camera’s field of view allowing us to record any scanning behaviours during the first few foraging trips, as the stereotypical route formed. After collecting ∼25 individuals’ outbound scans, we switched to focusing on recently emerged foragers on their inbound trip collecting any inbound scans, occurring near the nest. Once a forager completed a scan, or multiple scans within the recording frame, they were collected after they left the area and marked as completed to prevent repeated recording.

#### Experienced foragers procedure

Experienced forager scans dynamics were extracted and analysed from a data set published in Deeti et al. (2023), which focused specifically at scanning metric distributions; the following summary is further explained there. Two stocked feeders (cookie pieces) were sunk into the ground 5m from the nest, separated 120° (Right and Left sites). Ants exited each feeder via a 1m long, 10cm wide channel towards the nest, sloped up to ground level where ants exited onto a 60 cm × 120 cm ‘scanning platform’ (wooden board with white paper surface). Similar set-ups (minus the feeder) were placed at three other local sites (Middle, Opposite, and Far).

First, only the Right feeder was stocked with cookies and foragers were allowed to train along this route, with each visiting individual marked upon their first visit. Individuals were allowed to train along this route for two days prior to testing, repeatedly returning to the feeder and leaving via the channel with food. Once highly experienced, inbound foragers were tested by collection just before entering the nest and released at one of the four familiar test sites. As each individual reached the scanning platform they were video recorded using a high-speed camera (Casio EX-F1, FOV: 30×30 cm at 300 fps). This testing procedure was then repeated with a second group of foragers trained to the Left feeder. A final set up was placed in a far (40m), unfamiliar site where a separate group of experienced foragers was tested.

#### Pose extraction

For both sets of scan video datasets, we extracted the body orientations of ants during fixations to calculate their angle compared to the goal direction (e.g., nest or feeder) and their duration. Fixations were defined as periods when the ant’s head and body remained stationary between video frames, with no forward or rotational movement. Scanning can be composed of multiple scanning bouts, with little to no forward movement within a bout, while the ant rotates in place (saccades), or fixating in different directions, with a scanning bout ceasing once the ant resumes forward motion (Deeti et al., 2023a). By collecting body orientation during fixations, we extracted, the duration of each fixation, the ant’s *body orientation* relative to the goal, the angular change between each fixation (saccade angle) and duration, if the saccade was ‘towards’ or ‘away’ from the goal direction and when the fixation was followed by saccade that was a reversal in turning direction.

In the Experienced individuals (300 frames/s), orientation was extracted using a custom MATLAB code (see Deeti et al., 2023 for full description), with fixations identified as stationary periods between frames with body orientation estimated using points at the head and pronotum. In inexperienced forager scans (600frames/s), body orientation during fixations were determined manually using SLEAP software (Pereira et al., 2022). Fixations ‘started’ when the head stopped moving between frames and ended with the onset of either forward or rotational movement. Orientation was calculated using two body landmarks: the front centre of the head and the head/body connection (pronotum). As multiple scanning bouts can occur within a single path, each scanning bout began with the first, within frame fixation and ended after forward movement resumed for at least two full leg cycles. Additional fixations within this two-step window were still counted as within the same scanning bout.

### Statistical analysis (real-world Melophorus bagoti scans)

#### Grouping

For experienced foragers, scan data from all familiar release sites (Left, Right, Middle, Opposite) were pooled into a single familiar condition, while scans from the distant site were treated as unfamiliar. For inexperienced foragers, scans were separated into two groups based on if the scan occurred during the outbound and inbound portion of the foraging trip.

#### Linear mixed-effects models (LMEs)

LME models (Matlab2016) were used to assess the qualitative predictions of the model compared to our real ant data, rather than to optimize model fit. Accordingly, we report F-values derived from ANOVA tables, which test the significance of fixed effects rather than relying on t-values associated with individual parameter estimates. Importantly, this approach fits our qualitative focus. We first run models with interaction between factors. If no interaction was significant, we re-run the model for additive effect only.

Effects on *saccade aptitude* and *fixation duration* were analysed using Linear mixed-effect models with both *individual* and *test conditions* as random effects and with *angular deviation* from the goal direction as a continuous fixed effect and with reversal and Towards/Away (for fixations this corresponds with the upcoming saccade) as categorical fixed effects.

To assess the effect of experience on cumulative orientation vs. the goal during the first reversal of each scan, we conducted LME with the *cumulative orientation* as the dependent variable, with *condition* as the fixed effect and *individual* as a random effect to account for repeated measures across individuals. We ran a second LME to assess the effect of *experience* on the *saccade amplitude*, with *condition* as the fixed effect and *individual* as the random effect to account for repeated measures across individuals.

#### Fixation duration & saccade amplitude - linear regression

To assess the relationship between fixation duration and saccade amplitude, we first plotted the entire dataset. After observing the general trends, we separated shorter fixation durations (Quartile 1) from the rest of the dataset (Quartile 2-4) and performed separate linear regressions for both subsets. Regression slopes were then tested for significance using *t*-tests on the fitted model coefficients.

#### Start/stop anytime during oscillation cycle

To compare the duration via the number of saccades within a sweep between distinct sweeps (e.g., first vs. second sweep; penultimate vs. final sweep), we used the Wilcoxon signed-rank tests for paired data.

#### First reversal/longest fixation - ‘binomial’ chi square association

To assess if reversal and longest fixation duration were associated we tested them using the chi-square test of independence (χ²), and odds ratios with 95% confidence intervals were calculated.

## Supporting information

Vdeo 4

## Acknowledgements

We are grateful to the Centre for Appropriate Technology for permission to work on site and access to the nests. We thank Paul Graham for providing the experienced forager scan dataset, and Leo Clément and Gabriel G. Gattaux for contributing videos of ant behaviour.

## Funding Statement

The funders had no role in study design, data collection and interpretation, or the decision to submit the work for publication.

## Funding Information

This project was funded by a Macquarie University Research Fellowship (MQRF0001094) and by the European Research Council (RESIL-ANT - 101125881).

## Competing interests

No competing interests declared.

## Contributions

C.A.F. - Conceptualization, Data curation, Formal analysis, Funding acquisition, Validation, Investigation, Visualization, Methodology, Writing.

A.W. -Conceptualization, Data curation, Formal analysis, Modelling, Funding acquisition, Validation, Investigation, Visualization, Methodology, Writing.

## Data Availability Statement

All data, documentation and code is made available online at: (https://github.com/antnavteam/CX_oscillator_scans’)

## Notes

https://github.com/antnavteam/CX_oscillator_scans’

### Competing Interest Statement

The authors have declared no competing interest.

### Summary of Updates

Final Reviewer Comments for eLife had been added to this version (prerequisite of submitting revisions).

## References

Adden A, Stewart TC, Webb B, Heinze S. 2022. A Neural Model for Insect Steering Applied to Olfaction and Path Integration. Neural Comput 34:2205–2231. doi:10.1162/neco_a_01540

Baddeley B, Graham P, Husbands P, Philippides A. 2012. A Model of Ant Route Navigation Driven by Scene Familiarity. PLoS Comput Biol 8:e1002336. doi:10.1371/journal.pcbi.1002336

Baird E, Byrne MJ, Smolka J, Warrant EJ, Dacke M. 2012. The Dung Beetle Dance: An Orientation Behaviour? PLOS ONE 7:e30211. doi:10.1371/journal.pone.0030211

Beetz MJ, Kraus C, el Jundi B. 2023. Neural representation of goal direction in the monarch butterfly brain. Nat Commun 14:5859. doi:10.1038/s41467-023-41526-w

Berni J. 2015. Genetic Dissection of a Regionally Differentiated Network for Exploratory Behavior in *Drosophila* Larvae. Curr Biol 25:1319–1326. doi:10.1016/j.cub.2015.03.023

Berni J, Pulver SR, Griffith LC, Bate M. 2012. Autonomous Circuitry for Substrate Exploration in Freely Moving *Drosophila* Larvae. Curr Biol 22:1861–1870. doi:10.1016/j.cub.2012.07.048

Bidaye SS, Laturney M, Chang AK, Liu Y, Bockemühl T, Büschges A, Scott K. 2020. Two Brain Pathways Initiate Distinct Forward Walking Programs in *Drosophila*. Neuron 108:469–485.e8. doi:10.1016/j.neuron.2020.07.032

Buehlmann C, Fernandes ASD, Graham P. 2018. The interaction of path integration and terrestrial visual cues in navigating desert ants: what can we learn from path characteristics? J Exp Biol 221:jeb167304. doi:10.1242/jeb.167304

Buehlmann C, Mangan M, Graham P. 2020. Multimodal interactions in insect navigation. Anim Cogn 23:1129–1141. doi:10.1007/s10071-020-01383-2

Büschges A, Ache JM. 2025. Motor control on the move: from insights in insects to general mechanisms. Physiol Rev 105:975–1031. doi:10.1152/physrev.00009.2024

Cheng K. 2024. Oscillators and servomechanisms in navigation and orientation. Commun Integr Biol 17:2293268. doi:10.1080/19420889.2023.2293268

Cheng K, Narendra A, Sommer S, Wehner R. 2009. Traveling in clutter: navigation in the Central Australian desert ant Melophorus bagoti. Behav Processes 80:261–268.

Clément L, Schwarz S, Wystrach A. 2023. An intrinsic oscillator underlies visual navigation in ants. Curr Biol 33:411–422. doi:10.1016/j.cub.2022.11.059

Collett M. 2010. How desert ants use a visual landmark for guidance along a habitual route. Proc Natl Acad Sci USA 107:11638–11643. doi:10.1073/pnas.1001401107

Collett T, Collett M. 2002. Memory use in insect visual navigation. Nat Rev Neurosci 3:542–52. doi:10.1038/nrn872

Collett T, Graham P, Heinze S. 2025. The neuroethology of ant navigation. Curr Biol 35:R110–R124. doi:10.1016/j.cub.2024.12.034

Collett TS, Hempel De Ibarra N. 2023. An ‘instinct for learning’: the learning flights and walks of bees, wasps and ants from the 1850s to now. J Exp Biol 226:jeb245278. doi:10.1242/jeb.245278

Collett, TS, Hempel De Ibarra N, Riabinina O, Philippides A. 2013. Coordinating compass-based and nest-based flight directions during bumblebee learning and return flights. Journal of Experimental Biology, 216(6), 1105–1113.

Collett TS, Robert T, Frasnelli E, Philippides A, Hempel De Ibarra N. 2023. How bumblebees coordinate path integration and body orientation at the start of their first learning flight. J Exp Biol 226:jeb245271. doi:10.1242/jeb.245271

Dauzere-Peres O, Wystrach A. 2024. Ants integrate proprioception as well as visual context and efference copies to make robust predictions. Nat Commun 15:10205. doi:10.1038/s41467-024-53856-4

Dauzere-Peres O, De Wever S, Wystrach A. 2026. Predictive coding and oscillations underlie the optomotor response in distant insect lineages. bioRxiv, 2026–03.

Deeti S, Cheng K. 2025. Desert ants (*Melophorus bagoti*) oscillate and scan more in navigation when the visual scene changes. Anim Cogn 28:15. doi:10.1007/s10071-025-01936-3

Deeti S, Cheng K, Graham P, Wystrach A. 2023a. Scanning behaviour in ants: an interplay between random-rate processes and oscillators. J Comp Physiol A 209:625–639. doi:10.1007/s00359-023-01628-8

Deeti S, Islam M, Freas C, Murray T, Cheng K. 2023b. Intricacies of running a route without success in night-active bull ants (*Myrmecia midas*). J Exp Psychol Anim Learn Cogn 49:111–126. doi:10.1037/xan0000350

Demir M, Kadakia N, Anderson HD, Clark DA, Emonet T. 2020. Walking *Drosophila* navigate complex plumes using stochastic decisions biased by the timing of odor encounters. eLife 9:e57524. doi:10.7554/eLife.57524

Dudley R. 2000. The Biomechanics of Insect Flight: Form, Function, Evolution. Princeton University Press.

Fisher YE. 2022. Flexible navigational computations in the *Drosophila* central complex. Curr Opin Neurobiol, 73:102514.

Fleischmann PN, Grob R, Wehner R, Rössler W. 2017. Species-specific differences in the fine structure of learning walk elements in Cataglyphis ants. J Exp Biol 220:2426–2435. doi:10.1242/jeb.158147

Franconville R, Beron C, Jayaraman V. 2018. Building a functional connectome of the *Drosophila* central complex. eLife 7:e37017. doi:10.7554/eLife.37017

Freas CA, Cheng K. 2018. Landmark learning, cue conflict, and outbound view sequence in navigating desert ants. Journal of Experimental Psychology: Animal Learning and Cognition, 44(4): 409–421.

Freas CA, Cheng K. 2025. Visual learning, route formation and the choreography of looking back in desert ants, *Melophorus bagot*i. Anim Behav 222:123125.

Freas CA, Cheng K. 2022. The Basis of Navigation Across Species. Annu Rev Psychol 73:217–241. doi:10.1146/annurev-psych-020821-111311

Freas CA, Congdon JV, Plowes NJR, Spetch ML. 2020. Pheromone cue triggers switch between vectors in the desert harvest ant, Veromessor pergandei. Anim Cogn 23:1087–1105. doi:10.1007/s10071-020-01354-7

Freas CA, Fleischmann PN, Cheng K. 2019. Experimental ethology of learning in desert ants: Becoming expert navigators. Behav Processes 158:181–191. doi:10.1016/j.beproc.2018.12.001

Freas CA, Spetch ML. 2019. Terrestrial cue learning and retention during the outbound and inbound foraging trip in the desert ant, Cataglyphis velox. Journal of Comparative Physiology A. 205(2):177–89.

Freas CA, Spetch ML. 2023. Varieties of visual navigation in insects. Anim Cogn 26:319–342. doi:10.1007/s10071-022-01720-7

Freas CA, Whyte C, Cheng K. 2017. Skyline retention and retroactive interference in the navigating Australian desert ant, Melophorus bagoti. J Comp Physiol A Neuroethol Sens Neural Behav Physiol 203:353–367. doi:10.1007/s00359-017-1174-8

Freas CA, Wystrach A, Narendra A, Cheng K. 2018. The View from the Trees: Nocturnal Bull Ants, Myrmecia midas, Use the Surrounding Panorama While Descending from Trees. Front Psychol 9. doi:10.3389/fpsyg.2018.00016

Freas CA, Wystrach A, Schwarz S, Spetch ML. 2022. Aversive view memories and risk perception in navigating ants. Sci Rep 12:2899.

Green J, Maimon G. 2018. Building a heading signal from anatomically defined neuron types in the *Drosophila* central complex. Current opinion in neurobiology, 52, 156–164.

Gattaux G, Vimbert R, Wystrach A, Serres JR, Ruffier F. 2023. Antcar: Simple Route Following Task with Ants-Inspired Vision and Neural Model.

Gomez-Marin A, Louis M. 2014. Multilevel control of run orientation in *Drosophila* larval chemotaxis. Front Behav Neurosci 8. doi:10.3389/fnbeh.2014.00038

Gomez-Marin A, Stephens GJ, Louis M. 2011. Active sampling and decision making in *Drosophila* chemotaxis. Nat Commun 2:441. doi:10.1038/ncomms1455

Goulard, R, Buehlmann, C, Niven, JE, Graham, P, Webb, B. 2021. A unified mechanism for innate and learned visual landmark guidance in the insect central complex. PLoS computational biology, 17(9): e1009383.

Goulard R, Heinze S, Webb B. 2023. Emergent spatial goals in an integrative model of the insect central complex. PLOS Comput Biol 19:e1011480. doi:10.1371/journal.pcbi.1011480

Graham P, Collett TS. 2002. View-based navigation in insects: how wood ants (*Formica rufa* L.) look at and are guided by extended landmarks. J Exp Biol 205:2499–2509. doi:10.1242/jeb.205.16.2499

Green J, Vijayan V, Mussells Pires P, Adachi A, Maimon G. 2019. A neural heading estimate is compared with an internal goal to guide oriented navigation. Nat Neurosci 22:1460–1468. doi:10.1038/s41593-019-0444-x

Haalck L, Mangan M, Wystrach A, Clement L, Webb B, Risse B. 2023. CATER: Combined Animal Tracking & Environment Reconstruction. Sci Adv 9:eadg2094. doi:10.1126/sciadv.adg2094

Hangartner W. 1967. Spezifitat und Inaktivierung des Spurpheromons von Lasius fuliginosus Latr. und Orientierung der Arbeiterinnen im Duftfeld. Z Vgl Physiol 57:103–136. doi:10.1007/BF00303068

Heinze S. 2017. Unraveling the neural basis of insect navigation. Curr Opin Insect Sci 24:58–67. doi:10.1016/j.cois.2017.09.001

Honkanen A, Adden A, da Silva Freitas J, Heinze S. 2019. The insect central complex and the neural basis of navigational strategies. J Exp Biol 222:jeb188854. doi:10.1242/jeb.188854

Hulse BK, Haberkern H, Franconville R, Turner-Evans D, Takemura S, Wolff T, Noorman M, Dreher M, Dan C, Parekh R, Hermundstad AM, Rubin GM, Jayaraman V. 2021. A connectome of the *Drosophila* central complex reveals network motifs suitable for flexible navigation and context-dependent action selection. eLife 10:e66039. doi:10.7554/eLife.66039

Husbands P, Shim Y, Garvie M, Dewar A, Domcsek N, Graham P, Knight J, Nowotny T, Philippides A. 2021. Recent advances in evolutionary and bio-inspired adaptive robotics: Exploiting embodied dynamics. Appl Intell 51:6467 –6496. doi:10.1007/s10489-021-02275-9

Islam M, Freas CA, Cheng K. 2020. Effect of large visual changes on the navigation of the nocturnal bull ant, Myrmecia midas | Animal Cognition. Anim Cogn 23:1071–1080.

Iwano M, Hill ES, Mori A, Mishima Tatsuya, Mishima Tsuneko, Ito K, Kanzaki R. 2010. Neurons associated with the flip-flop activity in the lateral accessory lobe and ventral protocerebrum of the silkworm moth brain. J Comp Neurol 518:366–388. doi:10.1002/cne.22224

Izquierdo EJ, Lockery SR. 2010. Evolution and Analysis of Minimal Neural Circuits for Klinotaxis in *Caenorhabditis elegans*. J Neurosci 30:12908–12917. doi:10.1523/JNEUROSCI.2606-10.2010

Kanzaki R. 2005. Neural Basis of Odor-source Searching Behavior in Insect Brain Systems Evaluated with a Mobile Robot. Chem Senses 30:i285–i286. doi:10.1093/chemse/bjh226

Kanzaki, R., & Ikeda, A. (1994). Morphology and physiology of pheromone-triggered flip-flopping descending interneurons of the male silkworm moth, Bombyx mori. In Olfaction and Taste XI: Proceedings of the 11th International Symposium on Olfaction and Taste and of the 27th Japanese Symposium on Taste and Smell. Joint Meeting held at Kosei-nenkin Kaikan, Sapporo, Japan, July 12–16, 1993 (pp. 851–851). Tokyo: Springer Japan.

Kanzaki R, Mishima T. 1996. Pheromone-Triggered ‘Fiipflopping’ Neural Signals Correlate with Activities of Neck Motor Neurons of a Male Moth, Bombyx mori. Zoolog Sci 13:79–87. doi:10.2108/zsj.13.79

Kanzaki R, Nagasawa S, Shimoyama I. 2004. Neural Basis of Odor-Source Searching Behavior in insect Microbrain Systems Evaluated with a Mobile Robot In: Kato N, Ayers J, Morikawa H, editors. Bio-Mechanisms of Swimming and Flying. Tokyo: Springer Japan. pp. 155–170. doi:10.1007/978-4-431-53951-3_12

Kanzaki R, Sugi N, Shibuya T. 1992. Self-generated Zigzag Turning of Bombyx mori Males during Pheromone-mediated Upwind Walking(Physology). Zoolog Sci 9:515–527.

Kim SS, Hermundstad AM, Romani S, Abbott LF, Jayaraman V. 2019. Generation of stable heading representations in diverse visual scenes. Nature 576:126–131. doi:10.1038/s41586-019-1767-1

Knaden M, Graham P. 2016. The Sensory Ecology of Ant Navigation: From Natural Environments to Neural Mechanisms. Annu Rev Entomol 61:63–76. doi:10.1146/annurev-ento-010715-023703

Knight JC, Sakhapov D, Domcsek N, Dewar ADM, Graham P, Nowotny T, Philippides A. 2019. Insect-Inspired Visual Navigation On-Board an Autonomous Robot: Real-World Routes Encoded in a Single Layer NetworkThe 2019 Conference on Artificial Life. Presented at the The 2019 Conference on Artificial Life. Newcastle, United Kingdom: MIT Press. pp. 60–67. doi:10.1162/isal_a_00141

Kodzhabashev A, Mangan M. 2015. Route Following Without Scanning In: Wilson SP, Verschure PFMJ, Mura A, Prescott TJ, editors. Biomimetic and Biohybrid Systems. Cham: Springer International Publishing. pp. 199–210. doi:10.1007/978-3-319-22979-9_20

Kuenen LPS, Baker TC. 1983. A non-anemotactic mechanism used in pheromone source location by flying moths. Physiol Entomol 8:277–289. doi:10.1111/j.1365-3032.1983.tb00360.x

Legge, E. L., Wystrach, A., Spetch, M. L., & Cheng, K. (2014). Combining sky and earth: desert ants (Melophorus bagoti) show weighted integration of celestial and terrestrial cues. Journal of Experimental Biology, 217(23), 4159–4166.

Le Moël F, Stone T, Lihoreau M, Wystrach A, Webb B. 2019. The Central Complex as a Potential Substrate for Vector Based Navigation. Front Psychol 10:690. doi:10.3389/fpsyg.2019.00690

Le Möel F, Wystrach A. 2020. Opponent processes in visual memories: A model of attraction and repulsion in navigating insects’ mushroom bodies. PLOS Comput Biol 16:e1007631. doi:10.1371/journal.pcbi.1007631

Lehrer M. 1993. Why do bees turn back and look? J Comp Physiol A 172:549 –563. doi:10.1007/BF00213678

Lent DavidD, Graham P, Collett TS. 2010. Image-matching during ant navigation occurs through saccade-like body turns controlled by learned visual features. Proc Natl Acad Sci 107:16348–16353. doi:10.1073/pnas.1006021107

Li F, Lindsey JW, Marin EC, Otto N, Dreher M, Dempsey G, Stark I, Bates AS, Pleijzier MW, Schlegel P, Nern A, Takemura S, Eckstein N, Yang T, Francis A, Braun A, Parekh R, Costa M, Scheffer LK, Aso Y, Jefferis GS, Abbott LF, Litwin-Kumar A, Waddell S, Rubin GM. 2020. The connectome of the adult *Drosophila* mushroom body provides insights into function. eLife 9:e62576. doi:10.7554/eLife.62576

Lu J, Behbahani AH, Hamburg L, Westeinde EA, Dawson PM, Lyu C, Maimon G, Dickinson MH, Druckmann S, Wilson RI. 2022. Transforming representations of movement from body- to world-centric space. Nature 601:98–104. doi:10.1038/s41586-021-04191-x

Luo L, Gershow M, Rosenzweig M, Kang K, Fang-Yen C, Garrity PA, Samuel ADT. 2010. Navigational Decision Making in *Drosophila* Thermotaxis. J Neurosci 30:4261–4272. doi:10.1523/JNEUROSCI.4090-09.2010

Lyu C, Abbott LF, Maimon G. 2022. Building an allocentric travelling direction signal via vector computation. Nature 601:92–97. doi:10.1038/s41586-021-04067-0

Mishima T, Kanzaki R. 1999. Physiological and morphological characterization of olfactory descending interneurons of the male silkworm moth, Bombyx mori. J Comp Physiol [A] 184:143–160. doi:10.1007/s003590050314

Mouritsen H, Feenders G, Liedvogel M, Kropp W. 2004. Migratory Birds Use Head Scans to Detect the Direction of the Earth’s Magnetic Field. Curr Biol 14:1946–1949. doi:10.1016/j.cub.2004.10.025

Müller M, Wehner R. 2010. Path Integration Provides a Scaffold for Landmark Learning in Desert Ants. Curr Biol 20:1368–1371. doi:10.1016/j.cub.2010.06.035

Müller M, Wehner R. 2007. Wind and sky as compass cues in desert ant navigation. Naturwissenschaften 94:589–594. doi:10.1007/s00114-007-0232-4

Murakami M, Vicente MI, Costa GM, Mainen ZF. 2014. Neural antecedents of self-initiated actions in secondary motor cortex. Nat Neurosci 17:1574–1582. doi:10.1038/nn.3826

Murray T, Kocsi Z, Dahmen H, Narendra A, Le Möel F, Wystrach A, Zeil J. 2019. The role of attractive and repellent scene memories in ant homing (*Myrmecia croslandi*). J Exp Biol jeb.210021. doi:10.1242/jeb.210021

Muser B, Sommer S, Wolf H, Wehner R. 2005. Foraging ecology of the thermophilic Australian desert ant, Melophorus bagoti. Aust J Zool 53:301. doi:10.1071/ZO05023

Mussells Pires P, Zhang L, Parache V, Abbott LF, Maimon G. 2024. Converting an allocentric goal into an egocentric steering signal. Nature 626:808–818. doi:10.1038/s41586-023-07006-3

Namiki S, Iwabuchi S, Pansopha Kono P, Kanzaki R. 2014. Information flow through neural circuits for pheromone orientation. Nat Commun 5:5919. doi:10.1038/ncomms6919

Namiki S, Kanzaki R. 2016. The neurobiological basis of orientation in insects: insights from the silkmoth mating dance. Curr Opin Insect Sci 15:16–26. doi:10.1016/j.cois.2016.02.009

Nicholson DJ, Judd SPD, Cartwright BA, Collett TS. 1999. Learning walks and landmark guidance in wood ants (Formica rufa). J Exp Biol 202:1831–1838. doi:10.1242/jeb.202.13.1831

Olberg RM. 1983. Pheromone-triggered flip-flopping interneurons in the ventral nerve cord of the silkworm moth,Bombyx mori. J Comp Physiol A 152:297–307. doi:10.1007/BF00606236

Pereira TD, Tabris N, Matsliah A, Turner DM, Li J, Ravindranath S, Papadoyannis ES, Normand E, Deutsch DS, Wang ZY, McKenzie-Smith GC, Mitelut CC, Castro MD, D’Uva J, Kislin M, Sanes DH, Kocher SD, Wang SS-H, Falkner AL, Shaevitz JW, Murthy M. 2022. SLEAP: A deep learning system for multi-animal pose tracking. Nat Methods 19:486–495. doi:10.1038/s41592-022-01426-1

Pfeiffer K, Homberg U. 2014. Organization and Functional Roles of the Central Complex in the Insect Brain. Annu Rev Entomol 59:165–184. doi:10.1146/annurev-ento-011613-162031

Philippides, A, de Ibarra, NH, Riabinina, O, Collett, TS, 2013. Bumblebee calligraphy: the design and control of flight motifs in the learning and return flights of *Bombus terrestris*. Journal of Experimental Biology, 216(6), 1093–1104.

Rayshubskiy A, Holtz SL, Bates AS, Vanderbeck QX, Serratosa Capdevila L, Rockwell V, Wilson R. 2025. Neural circuit mechanisms for steering control in walking *Drosophila*. eLife 13:RP102230. doi:10.7554/eLife.102230.3

Robert T, Frasnelli E, De Ibarra NH, Collett TS. 2018. Variations on a theme: bumblebee learning flights from the nest and from flowers. J Exp Biol jeb.172601. doi:10.1242/jeb.172601

Sapkal N, Mancini N, Kumar DS, Spiller N, Murakami K, Vitelli G, Bargeron B, Maier K, Eichler K, Jefferis GSXE, Shiu PK, Sterne GR, Bidaye SS. 2024. Neural circuit mechanisms underlying context-specific halting in *Drosophila*. Nature 634:191–200. doi:10.1038/s41586-024-07854-7

Schwarz S, Clement L, Gkanias E, Wystrach A. 2020a. How do backward-walking ants (Cataglyphis velox) cope with navigational uncertainty? Anim Behav 164:133–142. doi:10.1016/j.anbehav.2020.04.006

Schwarz S, Mangan M, Webb B, Wystrach A. 2020b. Route-following ants respond to alterations of the view sequence. J Exp Biol 223:jeb218701. doi:10.1242/jeb.218701

Seelig JD, Jayaraman V. 2015. Neural dynamics for landmark orientation and angular path integration. Nature 521:186–191. doi:10.1038/nature14446

Shih C-T, Sporns O, Yuan S-L, Su T-S, Lin Y-J, Chuang C-C, Wang T-Y, Lo C-C, Greenspan RJ, Chiang A-S. 2015. Connectomics-Based Analysis of Information Flow in the *Drosophila* Brain. Curr Biol 25:1249–1258. doi:10.1016/j.cub.2015.03.021

Shiu PK, Sterne GR, Spiller N, Franconville R, Sandoval A, Zhou J, Simha N, Kang CH, Yu Seongbong, Kim JS, Dorkenwald S, Matsliah A, Schlegel P, Yu Szi-chieh, McKellar CE, Sterling A, Costa M, Eichler K, Bates AS, Eckstein N, Funke J, Jefferis GSXE, Murthy M, Bidaye SS, Hampel S, Seeds AM, Scott K. 2024. A *Drosophila* computational brain model reveals sensorimotor processing. Nature 634:210–219. doi:10.1038/s41586-024-07763-9

Steck K, Hansson BS, Knaden M. 2011. Desert ants benefit from combining visual and olfactory landmarks. J Exp Biol 214:1307–1312. doi:10.1242/jeb.053579

Steinbeck F, Adden A, Graham P. 2020. Connecting brain to behaviour: a role for general purpose steering circuits in insect orientation? J Exp Biol 223:jeb212332. doi:10.1242/jeb.212332

Stone T, Webb B, Adden A, Weddig NB, Honkanen A, Templin R, Wcislo W, Scimeca L, Warrant E, Heinze S. 2017. An Anatomically Constrained Model for Path Integration in the Bee Brain. Curr Biol 27:3069–3085.e11. doi:10.1016/j.cub.2017.08.052

Stürzl W, Zeil J, Boeddeker N, Hemmi JM. 2016. How Wasps Acquire and Use Views for Homing. Curr Biol 26:470–482. doi:10.1016/j.cub.2015.12.052

Sun X, Yue S, Mangan M. 2020. A decentralised neural model explaining optimal integration of navigational strategies in insects. eLife 9:e54026. doi:10.7554/eLife.54026

Tarsitano MS, Andrew R. 1999. Scanning and route selection in the jumping spider *Portia labiata*. Anim Behav 58:255–265. doi:10.1006/anbe.1999.1138

Tastekin I, Khandelwal A, Tadres D, Fessner ND, Truman JW, Zlatic M, Cardona A, Louis M. 2018. Sensorimotor pathway controlling stopping behavior during chemotaxis in the *Drosophila melanogaster* larva. eLife 7:e38740. doi:10.7554/eLife.38740

Ugolini A. 2006. Equatorial sandhoppers use body scans to detect the earth’s magnetic field. J Comp Physiol A 192:45–49. doi:10.1007/s00359-005-0046-9

Wallace GK. 1959. Visual Scanning in the Desert Locust Schistocerca Gregaria Forskål. J Exp Biol 36:512–525. doi:10.1242/jeb.36.3.512

Webb B, Wystrach A. 2016. Neural mechanisms of insect navigation. Curr Opin Insect Sci 15:27–39. doi:10.1016/j.cois.2016.02.011

Wehner R. 2009. The architecture of the desert ant’s navigational toolkit (Hymenoptera: Formicidae). Myrmecol News 12:85–96.

Wehner R, Fukushi T, Wehner S. 1992. Rotatory components of movement in high speed desert ants, Cataglyphis bombycina. Presented at the Proceedings of the 20th Göttingen Neurobiology Conference.

Wehner R, Hoinville T, Cruse H, Cheng K. 2016. Steering intermediate courses: desert ants combine information from various navigational routines. J Comp Physiol A 202:459–472. doi:10.1007/s00359-016-1094-z

Wehner R, Michel B, Antonsen P. 1996. Visual Navigation in Insects: Coupling of Egocentric and Geocentric Information. J Exp Biol 199:129–140. doi:10.1242/jeb.199.1.129

Wehner R, Srinivasan MV. 2003. Path Integration in Insects. Oxford University Press.

Westeinde EA, Kellogg E, Dawson PM, Lu J, Hamburg L, Midler B, Druckmann S, Wilson RI. 2024. Transforming a head direction signal into a goal-oriented steering command. Nature 626:819–826. doi:10.1038/s41586-024-07039-2

Wystrach A. 2023. Neurons from pre-motor areas to the Mushroom bodies can orchestrate latent visual learning in navigating insects. doi:10.1101/2023.03.09.531867

Wystrach A, Beugnon G, Cheng K. 2012. Ants might use different view-matching strategies on and off the route. J Exp Biol 215:44–55. doi:10.1242/jeb.059584

Wystrach A, Buehlmann C, Schwarz S, Cheng K, Graham P. 2020a. Rapid Aversive and Memory Trace Learning during Route Navigation in Desert Ants. Curr Biol 30:1927–1933.e2. doi:10.1016/j.cub.2020.02.082

Wystrach A, Lagogiannis K, Webb B. 2016. Continuous lateral oscillations as a core mechanism for taxis in *Drosophila* larvae. eLife 5:e15504. doi:10.7554/eLife.15504

Wystrach A, Mangan M, Philippides A, Graham P. 2013. Snapshots in ants? New interpretations of paradigmatic experiments. J Exp Biol jeb.082941. doi:10.1242/jeb.082941

Wystrach A, Mangan M, Webb B. 2015. Optimal cue integration in ants. Proc Biol Sci 282:20151484. doi:10.1098/rspb.2015.1484

Wystrach A, Moël FL, Clement L, Schwarz S. 2020b. A lateralised design for the interaction of visual memories and heading representations in navigating ants. doi:10.1101/2020.08.13.249193

Wystrach A, Philippides A, Aurejac A, Cheng K, Graham P. 2014. Visual scanning behaviours and their role in the navigation of the Australian desert ant Melophorus bagoti. J Comp Physiol A Neuroethol Sens Neural Behav Physiol 200:615–626. doi:10.1007/s00359-014-0900-8

Wystrach A, Schwarz S. 2013. Ants use a predictive mechanism to compensate for passive displacements by wind. Curr Biol 23:R1083–R1085. doi:10.1016/j.cub.2013.10.072

Yang HH, Brezovec BE, Serratosa Capdevila L, Vanderbeck QX, Adachi A, Mann RS, Wilson RI. 2024. Fine-grained descending control of steering in walking *Drosophila*. Cell 187:6290–6308.e27. doi:10.1016/j.cell.2024.08.033

Zeil J. 2023. Visual navigation: properties, acquisition and use of views. J Comp Physiol A 209:499–514. doi:10.1007/s00359-022-01599-2

Zeil J, Fleischmann PN. 2019. The learning walks of ants (Hymenoptera: Formicidae). Myrmecol News 29:93–110. doi:10.25849/MYRMECOL.NEWS_029:093

Zeil J, Kelber A, Voss R. 1996. Structure and function of learning flights in bees and wasps. Journal of experimental biology, 199(1), 245–252.

