## Supplemental for "Scanning and active sampling behaviours emerge from conserved insect neural circuits"

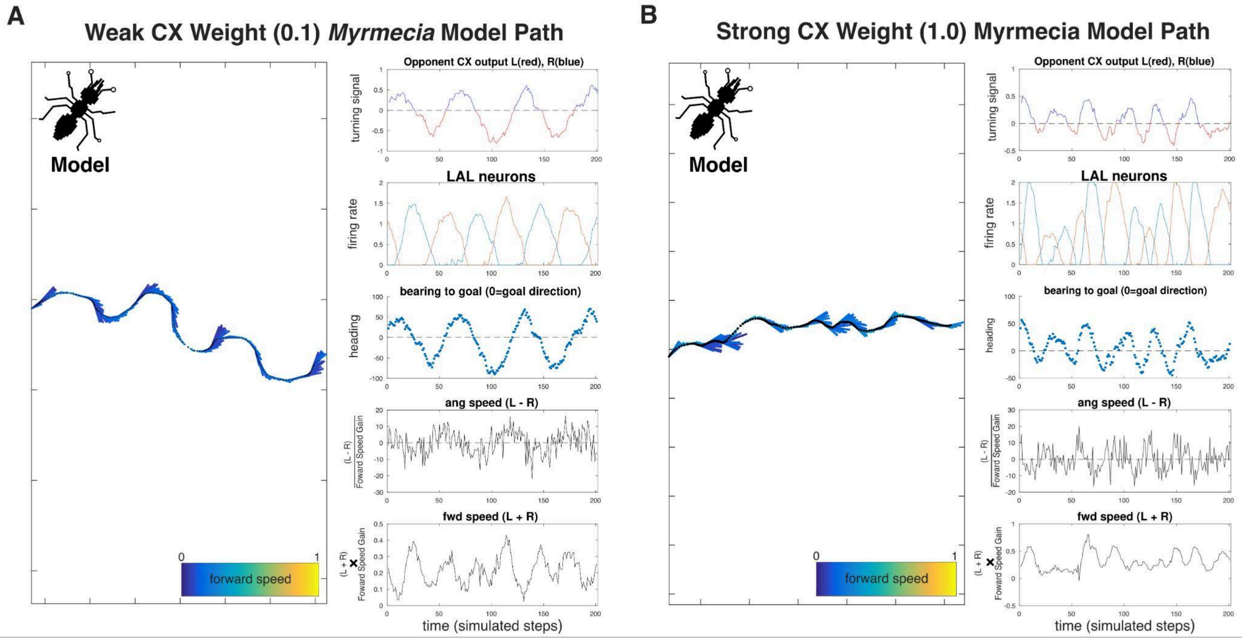


**Supplemental Figure 1. Two simulated agent paths under the '*Myrmecia*' model setting with a lower normal forward speed control range, with two settings for the CX signal weight.** (**A**) An example path under weak CX signal, CX signal = 0.1. (**B**) An example path under strong CX signal, CX = 1.0. Under stronger CX control, the oscillatory frequency increases (~1/40 steps) and is less regular compared to weaker CX steering signal weight (~1/60 steps). This is associated with path dynamics that are more goal directed and quicker in forward speed.

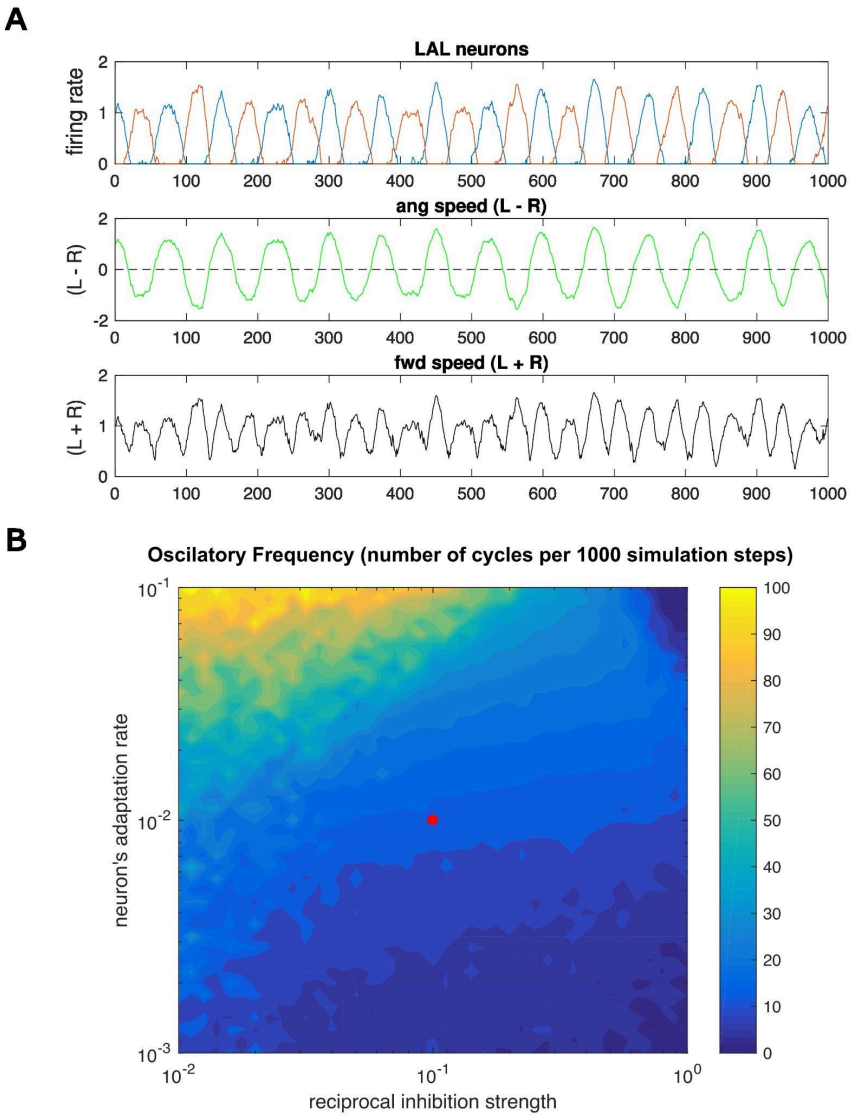


**Supplemental Figure 2. Dynamics of the oscillator across parameter ranges** A) Top Row: Firing rates of the two LAL neurons (red and blue) exhibiting antiphase oscillations. Middle Row: Angular speed (L - R), calculated as the difference between left and right motor outputs (green), driving turning behavior. Bottom Row: Forward speed (L + R), calculated as the sum of left and right motor outputs (black). (B) Heatmap showing oscillatory frequency as a function of reciprocal inhibition strength and neuronal adaptation rate. Color indicates the number of complete oscillatory cycles observed per 1000 simulation steps. Both axes are displayed on logarithmic scales. The red dot indicates the parameter combination used for the simulations shown in panel A and throughout the main figures. Parameter values were selected to ensure sufficient temporal resolution to capture multiple saccades within a single oscillatory cycle. Note that time is reported in dimensionless simulation steps rather than converted to milliseconds.
