## Supplementary material for "Scanning and active sampling behaviours emerge from conserved insect neural circuits": Video

**Video Captions**

**Video 1. Example of one of the full loop scans in inexperienced *Melophorus bagoti* foragers leaving the nest area towards the feeder**. Video was taken at 600fps at 1080p (~30cm x ~17cm) and is played at 60fps (slowed down 10x real time). The nest entrance is located ~50cm to the left of the frame while the goal is located 6m to the right of the frame (~90°). Video taken in Alice Springs, Northern Territory, Australia by CAF.

**Video 2. Illustrative example of a full loop scan in an outbound *Cataglyphis cursor* forager, along its foraging route in a familiar environment (experience level unknown).** Video taken by Gabriel G Gattaux in Marseille, France.

**Video 3. Illustrative example of a full loop scan/volte in a homing Myrmecia nigriceps navigating in an unfamiliar environment on a trackball. This video is provided as a qualitative example of the behaviour discussed in the text and was not included in the quantitative analyses*.*** Video taken in Sydney, New South Wales, Australia, by AW.

**Video 4. Illustrative example of oscillatory turning behaviour in a Myrmecia croslandi individual walking on a trackball while orienting in an unfamiliar environment. The video is provided as a qualitative example only and was not analysed quantitatively.** Video taken by Leo Clement in Canberra, Australian Capital Territory, Australia.

**Video 5. Video of example simulation of the modelled agent executing a full loop scan as depicted in Figure 7A.** Video shows fixations are in place and their sequence along with the corresponding CX and LAL outputs, the agents bearing and the angular (ang) and forward (fwd) speeds across time.

**Video 6. Video of example simulation of the modelled agent executing a double full loop scan as depicted in Figure 7B.** Video shows fixations are in place and their sequence along with the corresponding CX and LAL outputs, the agents bearing and the angular (ang) and forward (fwd) speeds across time.

**Video 7. Video of example simulation of the modelled agent executing a volte as depicted in Figure 7C.** Video shows fixations are in place and their sequence along with the corresponding CX and LAL outputs, the agents bearing and the angular (ang) and forward (fwd) speeds across time.
